# Starships are active eukaryotic transposable elements mobilized by a new family of tyrosine recombinases

**DOI:** 10.1101/2022.08.04.502770

**Authors:** Andrew S Urquhart, Aaron A Vogan, Donald M Gardiner, Alexander Idnurm

## Abstract

Transposable elements in eukaryotic organisms have historically been considered ‘selfish’, at best conferring indirect benefits to their host organisms. The *Starships* are a recently discovered feature in fungal genomes that are predicted to confer beneficial traits to their hosts and also have hallmarks of being transposable elements. Here, we provide experimental evidence that *Starships* are indeed autonomous transposons, using the model *Paecilomyces variotii*, and identify the HhpA ‘Captain’ tyrosine recombinase as essential for their mobilization into genomic sites with a specific target site consensus sequence. Furthermore, we identify multiple recent horizontal gene transfer of *Starships*, implying that they frequency jump between species. Fungal genomes have mechanisms to defend against mobile elements, which are frequently detrimental to the host. We discover that *Starships* are also vulnerable to repeat-induced point mutation defence, thereby having implications on the evolutionary stability of such elements.

## Introduction

Filamentous fungi of subphylum Pezizomycotina are one of the most speciose groups of organisms and have adapted to nearly every available ecological niche ^1,2^. Thus, is it perhaps not surprising that the genomes of these fungi are diverse and exhibit drastic shifts in gene content across relatively short evolutionary time spans ^3,4^. Mobile genetic elements are suspected agents of rapid adaptation in fungi, but the specific processes often remain obscure ^5^. In bacteria, mobile genetic elements can confer benefits to new hosts and thus facilitate adaptation to specific niches ^6,7^. Notably, Integrative and Conjugative Elements (ICEs) play a well-established role in disseminating the genetic information underlying adaptive traits. ICEs are mobile DNA (~20 kb to >500 kb in size) that contain the genes required for genomic integration, excision and transfer via conjugation ^8^. In addition, they contain a wide range of gene cargos conferring phenotypes such as antibiotic resistance, heavy metal resistance, nutrient utilisation, and pathogenicity (reviewed by ^8,9^).

Until now a parallel role in carrying host-beneficial DNA has not been established for mobile elements in Eukaryotes - despite mobile elements having first been discovered in, and being abundant in, many eukaryotic genomes. For example, in humans almost half (45%) of the genome consists of remnant transposable elements (TEs), and one element alone, Alu, is present in over 1 million copies ^10,11^. Mobile elements in eukaryotes have historically been considered selfish pieces of DNA and in fungi much research has focused on how the genome defends itself against their deleterious impacts ^12^. More recently, an appreciation of the role of TEs in eukaryotic adaptation has emerged but is focused on inadvertent effects such as insertional mutation, altered gene expression in neighbouring genes, and genome instability ^13^. For example, a transposon inserted within the gene *cortex* was responsible for increased melanisation in peppered moths (*Biston betularia*) in response to pollution during the Industrial Revolution ^14^. Similarly, resistance to azole fungicides in many species of fungi results from insertion of transposons into the *CYP51* gene promoter ^15–18^. In both these situations the gene products of the transposons is not inherently beneficial to the host but happens to be inserted at a particular location which influences the expression of adjacent genes. Examples of eukaryotic transposons that directly benefit the new host are sparse ^19^. However, one example of inherently host-beneficial transposons is known from the plant pathogen *Botrytis cinerea* wherein the fungal retrotransposons manipulate plant gene expression by encoding trans-species sRNAs ^20^.

Recently, several large genomic elements have been discovered in fungi that appear to carry protein-encoding host-beneficial genes in the way that prokaryotic ICE elements do. One such element, *HEPHAESTUS* (*Hφ*), was discovered in *Paecilomyces variotii*, which is a member of the order Eurotiales that also includes the better-known genera *Aspergillus* and *Penicillium. Hφ* carries a diverse set of cargo genes that provide resistance to at least five metal ions: zinc, cadmium, arsenic, copper, and lead. Additionally, *Hφ* appears to have undergone horizontal gene transfer (HGT) to *P. variotii* from *Penicillium fuscoglaucum* or a closely related species. Related elements appear to be common among fungi in the Eurotiales order, including multiple species of *Penicillium* and *Aspergillus*, carrying a diverse range of different genes putatively involved in metal ion tolerance, suggesting that these element are widespread, yet polymorphic in their gene content, among these fungi ^21^.

*Hφ* and a multitude of structurally similar elements have now been categorized as belonging to a novel class of genetic elements called ‘*Starships*’, which bioinformatic searches reveal to be abundant within Pezizomycotina genomes ^22^. The *Starships* encompass large (up to ~300 kb in *Galactica*) genomic regions that exhibit characteristics consistent with being mobile elements. Firstly, they frequently show presence/absence polymorphisms within species, and secondly, they vary in their position within the genome of different strains ^21,22^.

The most frequently found gene in *Starships* is a putative tyrosine recombinase (YR), currently assigned to the pfam classification domain of unknown function (DUF) 3435 ^22,23^. In *Hφ* the homolog has been named HhpA ^21^. Homologs of this gene are always found in the first position in *Starships*, and like canonical YRs, possesses a conserved catalytic pentad RKHRY ^24^. It is hypothesized that this gene alone is responsible for the mobilization of the *Starships*, but this has yet to be demonstrated ^24^. Additionally, no *Starship* has been observed to actively translocate, neither within nor between genomes.

In addition to the experimentally characterized *Hφ* example, predicted *Starships* contain diverse genes including biosynthetic gene clusters and putative pathogenicity determinants, much of which may be similarly host-beneficial ^22^. In at least one case, a *Starship* contains meiotic drive genes, which allow it to bias its transmission to offspring ^24^. It has thus been proposed that these *Starships* could represent eukaryotic analogues of prokaryotic ICEs ^21,22^. A striking parallel between *Starships* and bacterial ICEs is that they often possess signatures of HGT. The vast number of sequenced genomes has led to the realization of recurrent HGT within fungi ^25^. Furthermore, these transfers are not limited to individual genes, but often encompass entire gene clusters ^26^. Many of these cases have since been shown to involve *Starship* elements ^20^, uniting many disparate observations under a single phenomenon, and implicating *Starships* as vehicles of HGT in fungi. While bioinformatic support for such roles is strong, key questions have until now remained unanswered. (1) Are the *Starships* indeed actively mobile within genomes? (2) If so, by what mechanism do they move? (3) Do these play an analogous role to prokaryotic ICEs in moving diverse genetic cargo horizontally between species? (4) How do *Starships* interact with the host genome defences?

## Results and Discussion

### A *Starship*, HEPHAESTUS, is actively mobile within the fungal genome

We focused on the *Starship* called *HEPHAESTUS* (*Hφ*) of the fungus *Paecilomyces variotii*. This is because it is found within an experimentally tractable organism ^27^, contains a relatively well characterised set of host-beneficial genes conferring tolerance to metal ions at toxic concentrations, and near identical copies are variably located in different strains suggesting it may be actively transposing ^21^.

To observe *Hφ* movement we created a reporter strain for movement by disrupting an antibiotic resistance cassette with *Hφ* positioned between the promoter and coding sequence for hygromycin phosphotransferase (see illustration in Fig. 1A, 1B). Movement of *Hφ* by excision resulted in hygromycin resistant colonies at a frequency of approximately 1 excision per 10^6^ to 10^7^ spores. As the *Hφ* element carries genes for zinc and cadmium resistance (conferred by the *zrcA* and *pcaA* genes ^21^), the hygromycin resistant colonies were screened on media containing high concentrations of these metal ions. Resistance has been maintained in 13/15 colonies, indicating that *Hφ* has reintegrated elsewhere within the genome of these strains (Fig. 1C). To identify the reintegration positions of *Hφ*, these 13 colonies were whole genome sequenced as a single pool. Seven independent insertion sites were identified within this pool. These confirmed that *Hφ* preferentially integrates into genomic TTAC sites, in agreement with previously reported data on naturally occurring *Hφ* copies ^21^ (Fig. 1D). Sequencing of the excision sites revealed no remnant DNA from *Hφ* at the excision site (Fig. 1E), suggesting that the repeats found on the edges of the *Starships* are not derived through duplication of the target site ^24^; rather, we hypothesize that the elements contain sequence homologous to the target site at their termini.

**Fig. 1.**
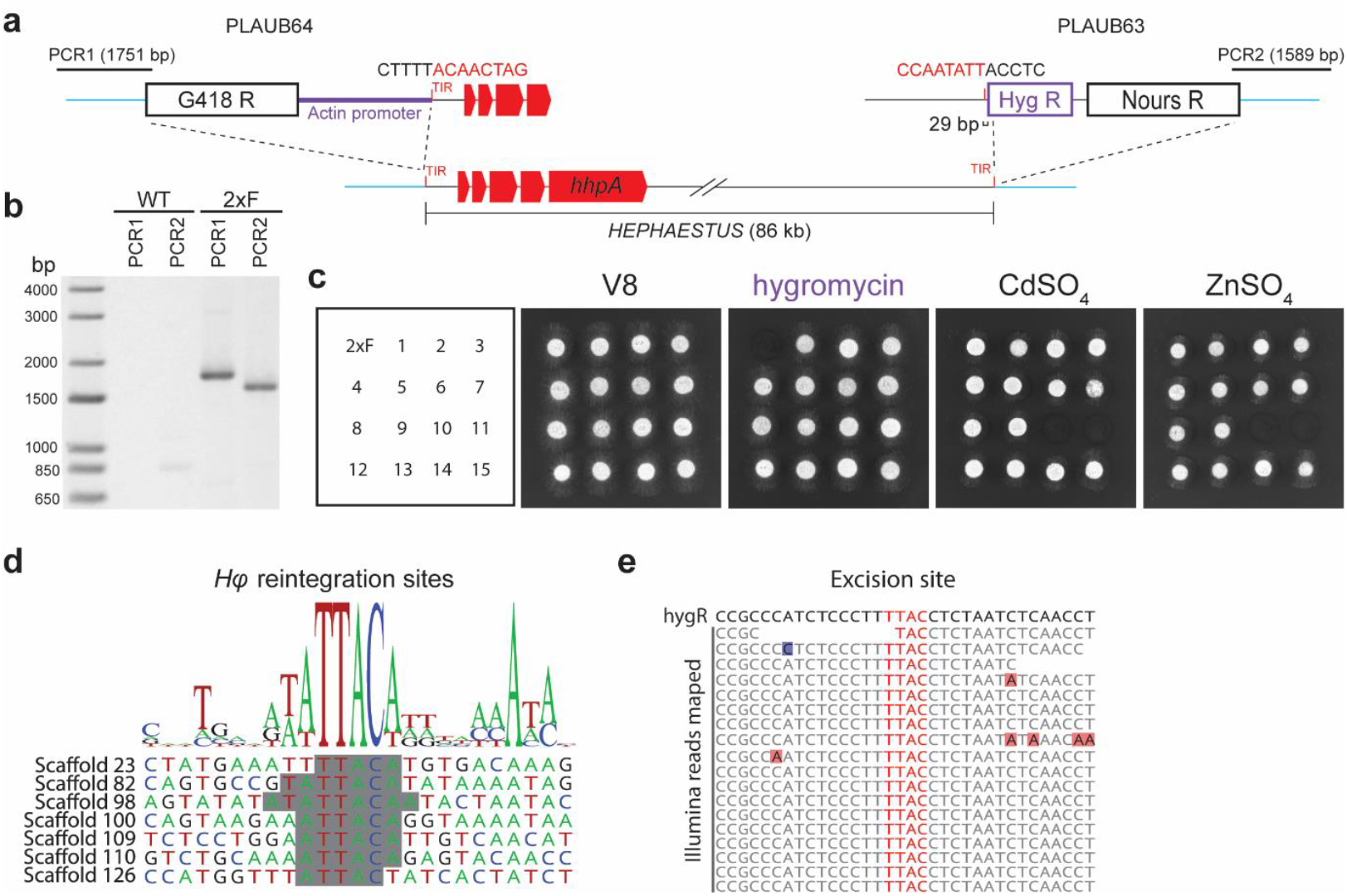
*HEPHAESTUS* is actively mobile within the *Paecilomyces variotii* genome. **a**, Two constructs were transformed on either side of *Hφ* within the CBS 144490 genome. This arrangement leads to the expression of hygR only after spontaneous excision of the *Hφ* region because this brings the *Leptosphaeria* actin promoter and hygromycin phosphotransferase coding region together. **b**, PCR analysis confirmed the correct integration of PLAUB64 and PLAUB63 either side of *Hφ*. **c**, Growth of a “double flanked” transformant (2xF) strain and spontaneously hygromycin resistance derivatives cultured on medium supplemented with hygromycin, cadmium and zinc. **d**, Reintegration sites of *Hφ* within the genome identified via pooled Illumina sequencing. Shading in alignments presents the direct DNA repeats created due to matching DNA between *Hφ* termini and target sites. **e**, Illumina sequencing reads mapped to the hygR cassette revealed no footprint left at the *Hφ* excision site.

### A putative tyrosine recombinase, the ‘Captain’, drives the movement of *HEPHAESTUS*

Having demonstrated that *Hφ* can move within the genome we next sought to understand how this movement occurs. A feature common to all *Starships* is the presence of a gene homologous to *hhpA* that have been termed ‘Captains’ given that they ‘lead’ the cluster ^22,24^. These captain proteins show weak similarity to tyrosine recombinase (YR) enzymes, which are known to mobilise other mobile elements such a *Cryptons* ^28^ and are the sole protein required for integration of other mobile elements such as the *Saccharomyces cerevisiae* 2µ plasmid ^29,30^.

As evidence that HhpA is a recombinase we investigated its subcellular localisation, expected to be nuclear in the case of a protein that interacts with DNA. We therefore made a strain expressing a fusion of HhpA to GFP (C-terminal tag at native locus) and examined fluorescence in the cell. Spherical sites of fluorescence were observed, consistent with nuclei. To confirm this, the strain was transformed with a construct expressing a histone H2B-mScarlet fusion, which exhibited co-localization between the red and green signals (Fig. 2A).

**Fig. 2.**
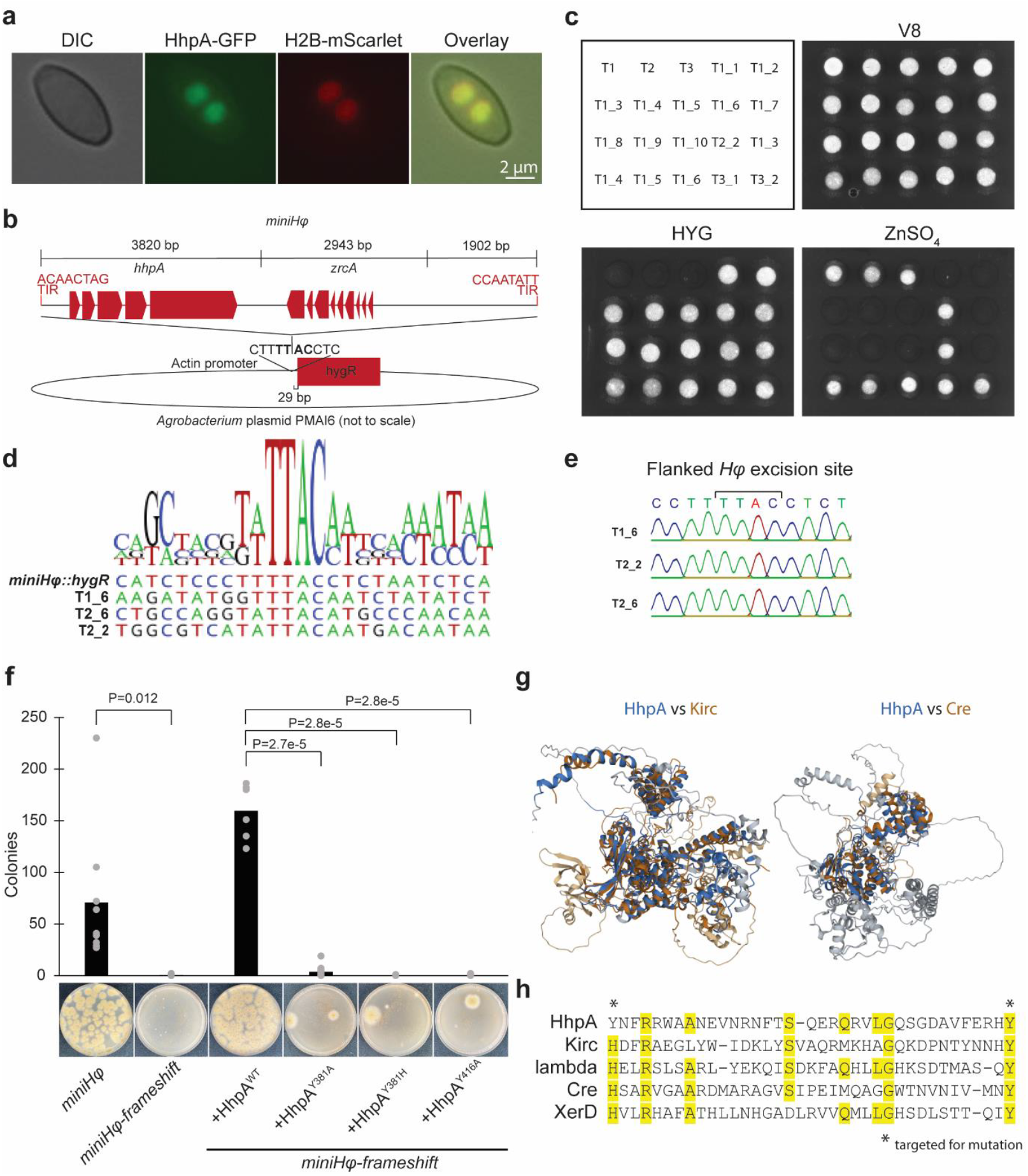
*HEPHAESTUS* movement is mediated by the putative tyrosine recombinase enzyme HhpA. **a**, Nuclear localisation of a HhpA-GFP fusion protein. **b**, A minimal version of *Hφ* was inserted into plasmid PMAI6 29 bp upstream of the hygromycin phosphotransferase coding region. Spontaneous excision of *Hφ* will result in expression of the *hygR* gene conferring hygromycin resistance. **c**, Growth of *miniHφ* transformants and their spontaneously hygromycin resistant derivatives on hygromycin and zinc. **d**, Reintegration sites of the *miniHφ* element determine using inverse PCR. **e**, Sanger sequencing of excision site indicates shows that no DNA (ie footprint) remains. **f**, AlphaFold protein structure predictions for HhpA and Kirc show structural similarity despite the low level of amino acid sequence similarity between these proteins. The HphA predicted structure also shows similarity to the Cre recombinase crystal structure (PDB: 3CRX.E) **g**, Sequence alignment of HhpA at part of the putative tyrosine recombinase active site previously identified by Vogan et al. **h**, Frameshift mutation of HhpA reduced transposition frequency which could be restored by WT HhpA but not by HhpA Y381A, Y381H or Y416A.

To experimentally demonstrate a role for HhpA in the mobilisation of *Hφ*, a minimal version of *Hφ* (“*miniHφ*”) was created, that features DNA from either flank of *Hφ* (5’
s 3820 bp, 3′ 1902 bp), the *hhpA* gene, and the *zrcA* gene. This version was inserted within the promoter driving hygromycin resistance (illustrated in Fig. 2B). The *zrcA* gene allows selection of zinc resistant transformants when introduced into a strain lacking a naturally occurring copy of *zrcA* (carried within *Hφ*). Like the approach with full length *Hφ* (Fig. 1A), the initial transformants selected on zinc were sensitive to hygromycin, and therefore to identify events in which *miniHφ* was excised, conidia collected from strains grown on the non-selective media were plated onto medium containing hygromycin. Some of these hygromycin resistant strains had reintegrated the *miniHφ* elsewhere in the genome, as expected if transposition had occurred (Fig. 2C and D). As with the full length *Hφ*, amplification, sequencing and analysis of the DNA revealed that no footprint was left behind following excision (Fig. 2E). To confirm the role of the HhpA protein product in the movement of *miniHφ*, a similar construct with a frameshift mutation in the *hhpA* gene was generated, and then subjected to the same approach of transformation with selection using zinc resistance, and screening for element excision by growth on hygromycin. This mutation resulted in a greatly reduced excision frequency, indicating that the protein is required for successful transposition (Fig. 2F). Complementation of the strain *miniHφ-frameshift* with a construct expressing HhpA under the control of a constitutive promoter restored excision frequency, and demonstrated that the transposition function can be provided *in trans*.

Vogan et al. ^24^ showed that the ‘Captain’ protein, KIRC has some similarity to YRs, but the protein sequence identity between KIRC and HhpA is low (~14%). To further investigate the Captain proteins we computationally predicted structures using AlphaFold ^31^. The predictions showed considerable similarity for KIRC and HhpA (HhpA aa 8 to 671, KIRC aa 1 to aa 584, TM-score 0.71) as well as between HhpA and Cre recombinase (HhpA aa 39 to 431, KIRC aa 33 to aa 341, TM-score 0.59) (Fig 2G). This strengthens the evidence that these two captain proteins share a common function and that they both represent tyrosine recombinase enzymes. Interestingly, while KIRC possess a histidine residue in the active site, concordant with most other YRs, HhpA has a tyrosine at this position (Y381) (Fig. 2H). Surveying the broader phylogeny of Captain protein sequences reveals that this polymorphism occurs in a large number of Captains, and appears to be an ancient division, bisecting a phylogeny of the Captains at a deep node (Fig. S1). To test whether HhpA is a YR, we mutated key residues in the predicted tyrosine recombinase active site (Fig. 2H), namely Y381 and Y416, within the construct used above to complement the *miniHφ-frameshift* strain. We then transformed these into *miniHφ*-*frameshift* and tested if these could restore transposition frequency in the *miniHφ*-*frameshift* strain as the wild type HhpA was shown to do: mutation of these residues to an alanine resulted in a decrease in the frequency of excision (Fig. 2F) further supporting their identity as YR enzymes. Furthermore, mutation of Y381 to a histidine was also non-functional (Fig. 2F) consistent with a deep divergence among the Captain lineages.

Exactly how the Captain proteins mediate movement of *Starships* is unclear but one possibility is through a circular intermediate as hypothesised, but not yet demonstrated, for *Crypton* transposons ^28^.

### *HEPHAESTUS* preferentially inserts at a specific target site

Based on examination of a limited number of natural *Hφ* insertion sites we previously identified a preference towards insertion into TTAC sites ^21^. YR enzymes frequently require greater regions of homology, for example the biotechnologically important Cre recombinase is specific to the 34 bp *loxP* sequence ^32^. As such we decided to generate a larger set of *Hφ* target sites to search for additional conservation at the integration sites of *Hφ*. To do this we collected a large number of strains in which *Hφ* had jumped (using hygromycin selection as described previously) and sequenced these as a pool. Reads covering the flanks of *Hφ* were extracted from the resultant data via mapping to the termini of *Hφ* and the genomic DNA flanking the *miniHφ* sequence was mapped back onto the genome to identify *Hφ* insertions. In total 132 insertion sites were identified in which both genomic flanks were unambiguously mapped (Fig 3A; Fig. S2). Examining the larger number of insertions shows that there is additional conservation around the previously identified TTAC site. Most strikingly there is a highly conserved adenosine (98%) downstream from the TTAC site at position +10 (Fig. 3B, Table S1).

**Fig. 3.**
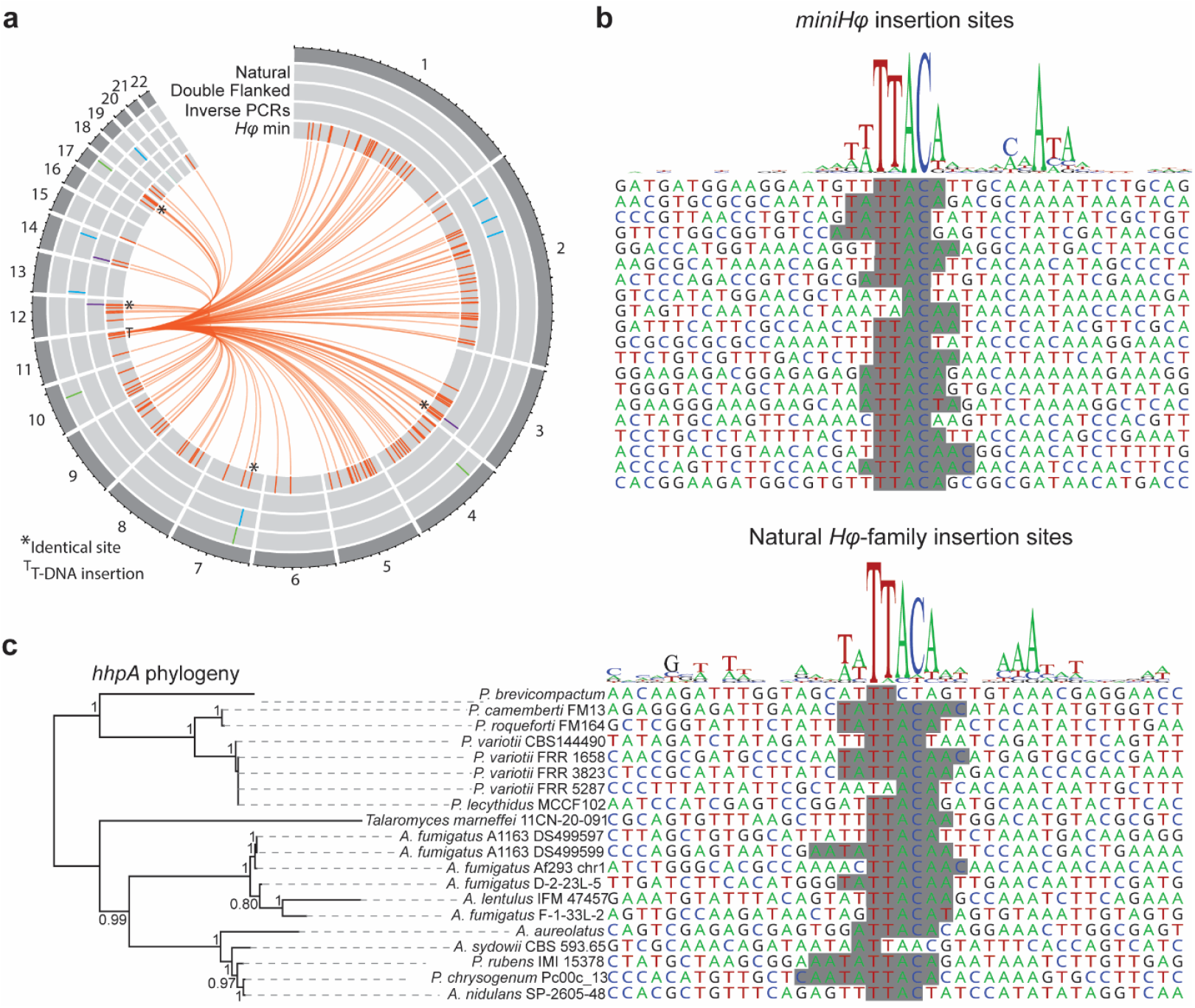
*HEPHAESTUS* integrates into TTAC(N_7_)A sites. **a**, Position of the four known natural insertions, seven experimentally isolated insertions, the three *miniHφ* insertion identified by inverse PCR and the 132 *miniHφ* insertion sites from pooled Illumina reads on the 22 largest *P. variotii* CBS 101075 assembly scaffolds. Internal lines connect the 132 *miniHφ* insertion sites to the location of the T-DNA on which the *miniHφ* sequence was introduced. **b**, Consensus sequence of all 132 *miniHφ* target sites, and an alignment of a subset of 20 sites from 132 characterised (a complete extended alignment is given in Fig. S2). **c**, Consensus sequence at the insertion site of naturally occurring *Hφ* relatives. Insertion sites are ordered according to the phylogenetic relationships of the associated *HhpA* homologs. Shading in alignments presents the direct DNA repeats created due to matching DNA between *Hφ* termini and target sites.

To determine whether these target site features are likely to be a conserved feature of the *Hφ-family* of *Starships* we compared the insertion sites of *miniHφ* to the insertion sites of naturally occurring *Hφ* homologs (Fig. 3C, Table S1). Twenty such *Starship* sites were either observed previously ^21^ or are newly identified *Hφ* relatives that could be cleanly define based on comparison to a corresponding empty site (Table S2). The features described above are generally conserved among *Hφ*-family members. Target site preference does not appear to vary between different *Hφ*-family starships despite the evolutionary distance between their Captains proteins.

These results indicate that the preferred target site is likely conserved within *Starship* families. They also indicate that *Starship* target sites might be more complex than could be identified from a small number of insertion sites ^21,24^ and should thus be interpreted in the context of multiple insertion site alignments where possible. The integration of ICE elements into the host chromosome is also frequently mediated by tyrosine recombinases. ICE elements show a considerable diversity in their specificity for a particular integration site, ranging from integration into a single att site within the genome to non-specific integration as in Tn*916* ^8,33^. This diversity appears to be mirrored within the *Starships* with some such as *Hφ* having relatively low specificity, with at least hundreds of potential sites in the genome and a relatively simple consensus sequence, while others such as *Voyager* show greater specificity – in this case to the 5S rDNA ^22^.

### The complete *HEPHAESTUS* has been horizontally transferred between two *Paecilomyces* species

We previously uncovered signal of horizontal transfer for blocks of genes contained within *Hφ*-family *Starship* elements ^21^. This was highly suggestive of a role for *Hφ*-family *Starships* in transferring genes between species. However, despite this strong evidence, an observation of two near-identical full length *Hφ* elements in different species remained elusive ^21^. Further bioinformatic searches of recently released GenBank sequences identified an example in *Paecilomyces lecythidis* strain MCCF 102 (Fig. 4A). This genome contains a full length *Starship* that is near-identical to *Hφ* of *P. variotii* (only one nucleotide mutated in 86 Kbp of sequence which was not attributable to sequencing error).

**Fig. 4.**
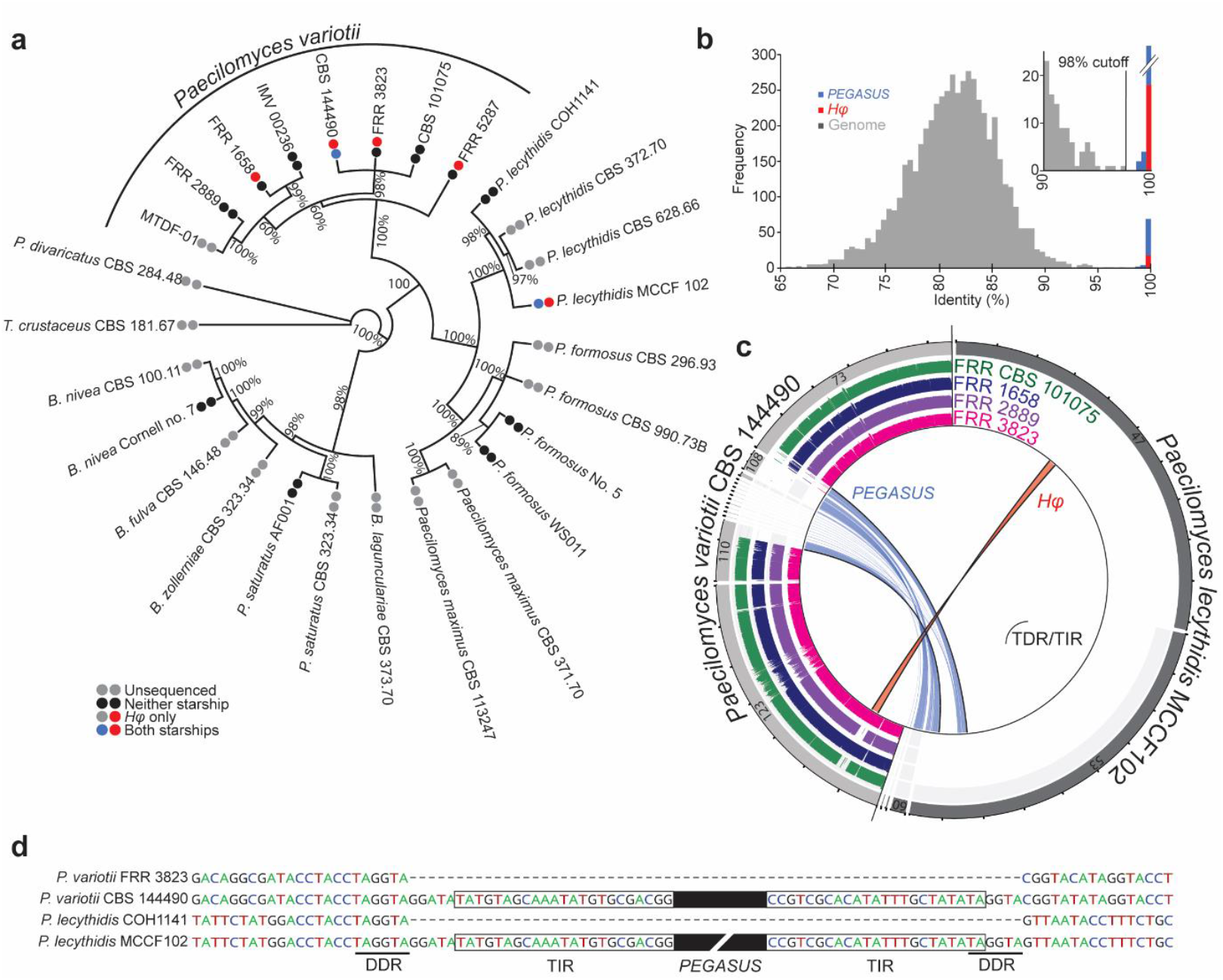
Evidence of *HEPHAESTUS* transfer between fungal species. **a**, Phylogenetic tree based on the calmodulin marker showing the taxonomic relationships between publicly available *Paecilomyces* genomes and taxonomically certain strains from ^35^. Tree generated in MrBayes ^36^ branch supports represent posterior probabilities. Distribution of *Hφ* and a second *Starship PEGASUS* is indicated by red and blue dots, respectively. **b**, A histogram representing the top BLASTn search result for each 5 kb window of the *P. variotii* CBS 144490 genome against the *P. lecythidis* MCCF 102 contigs. The presence of highly conserved genomic regions is indicative of HGT. A cut-off of 98% was used to predict putative HGT. **c**, A Circos plot showing the genomic location all BLASTn matches with >98% identity as identified in panel B. Lines drawn between the location of the BLAST queries on the *P. variotii* CBS 144490 contigs and subject sequences on the *P. lecythidis* MCCF 102 contigs. Red indicates the *Hφ* region and blue represents a second *Starship PEGASUS* that has also undergone HGT, albeit is fragmented in both assemblies. *PEGASUS* of MCCF 102 has undergone an inversion mutation since it was integrated into the genome as described in Fig. S3. **d**, Nucleotide sequence alignment of the edges of *PEGASUS* in *P. variotii* 144490 and *P. lecythidis* COH1141 compared to the corresponding empty sites in *P. variotii* FRR 3823 and *P. lecythidis* MCCF 102. Key *Starship* features were present including a terminal inverted repeat (TIR) and a direct DNA repeat (DDR).

To further illustrate that this represents a HGT event between these species we conducted a “BLAST all” analysis similar to what we have described previously ^21^. This approach was taken to avoid vulnerabilities associated with phylogenetic approaches such as introgression or incomplete lineage sorting ^34^. Briefly, we fragmented the *P. variotii* genome into 5 Kbp fragments and used BLASTn to compare these against the *P. lecythidis* MCCF 102 contigs. We then produced a histogram representing the frequency of various percentage identities in the resulting BLAST hits (excluding those less than 100 bp in length). This showed that the *Hφ* region (in red) is more conserved than the bulk of the genome (in grey) which has presumably been vertically inherited from a common ancestor (Fig 4B), providing strong evidence for horizontal transfer.

Unexpectedly, other genomic fragments (in blue) also showed high conservation (Fig 4B). Mapping all fragments with conservation >98% back onto the CBS 144490 contigs revealed four main regions of high conservation, one of which was *Hφ* (Fig. 4C). These potentially represented a second *Starship* that is fragmented in the genome assemblies, which we name *PEGASUS* after the Greek mythical winged horse. To demonstrate that these regions represent a single element we confirmed the co-segregation of these regions in 22 progeny of a cross between a *PEGASUS* + strain and *PEGASUS* - strain of *P. variotii* (Table S3). Examination of the sequence revealed TIR and direct DNA repeats (DDR) characteristic of *Starships* (Fig. 4D). Furthermore, contig 110 contains a gene encoding a “Captain” recombinase homolog, *pgcA* “*PEGASUS* captain A”, which has a DUF3435 domain. It thus appears that at least two *Starships* have undergone HGT between *P. variotii* and *P. lecythidis*. Further compelling evidence that *PEGASUS* is a recent transfer between genomes is its rapid mutation that occurs during sexual reproduction, which will be described in a following section.

*Starships*, in addition to encoding ‘Captains’ like HhpA, frequently encode a number of other auxiliary genes. These include a gene with a DUF3723 domain, a putative iron transporter (FreB), a gene with a patatin-like phospholipase domain, and a variety of other genes that can be classified as NOD-like receptors (NLRs) ^22^. It has been hypothesized that one or more of these additional genes might be involved in horizontal gene transfer ^22^. The DUF3723 domain has distant homology to bacterial proteins that stabilize chromosomes during division, and NLRs often mediate hyphal fusion in fungi, which could provide the means for strain to strain transfer in an analogous process to bacterial conjugation ^22^. *Hφ* lacks a functional homolog of any of these genes (although related metal resistance *Starships* do have them), yet has clearly undergone HGT suggesting that these genes are not required for *Hφ* movement. We cannot exclude the possibility that a second element such as *PEGASUS* has provided *in trans* the required auxiliary genes for the transfer of *Hφ*, but it is at least clear that the entire *Hφ* sequence can move between species without the presence of these genes within it. The mechanisms of *Starship*-mediated HGT are a major frontier for future research.

### *HEPHAESTUS* elements carry diverse host-beneficial cargo between fungal genomes

While we have described a number of *Hφ*-family elements that contain metal resistance genes ^21^, more broadly, *Starships* contain a much wider range of potentially host-beneficial gene content ^22^. Thus, we hypothesized that *Starship*-mediated HGT (including by the *Hφ*-family) may be a broader phenomenon involved in a wider range of phenotypes. Unfortunately, few *Starships* are yet sufficiently well-characterised to infer an impact on the host’s physiology or to be implicated in a HGT event. The exceptions are *Hφ* and the *Horizon* carrying the pathogenicity factor ToxA ^21,37^. To understand if the *Hφ*-family HGT events are specific to metal resistance we first identified a set of full length *Hφ* elements (Fig. S4, Table S4). These were chosen using a gene-content neutral approach. Several of these contain no putative metal resistance genes, but instead possess genes with other predicted cellular functions. Among these elements we have identified two cases of HGT of different host-beneficial cargo (Fig. 5A, Fig. 5B).

**Fig. 5.**
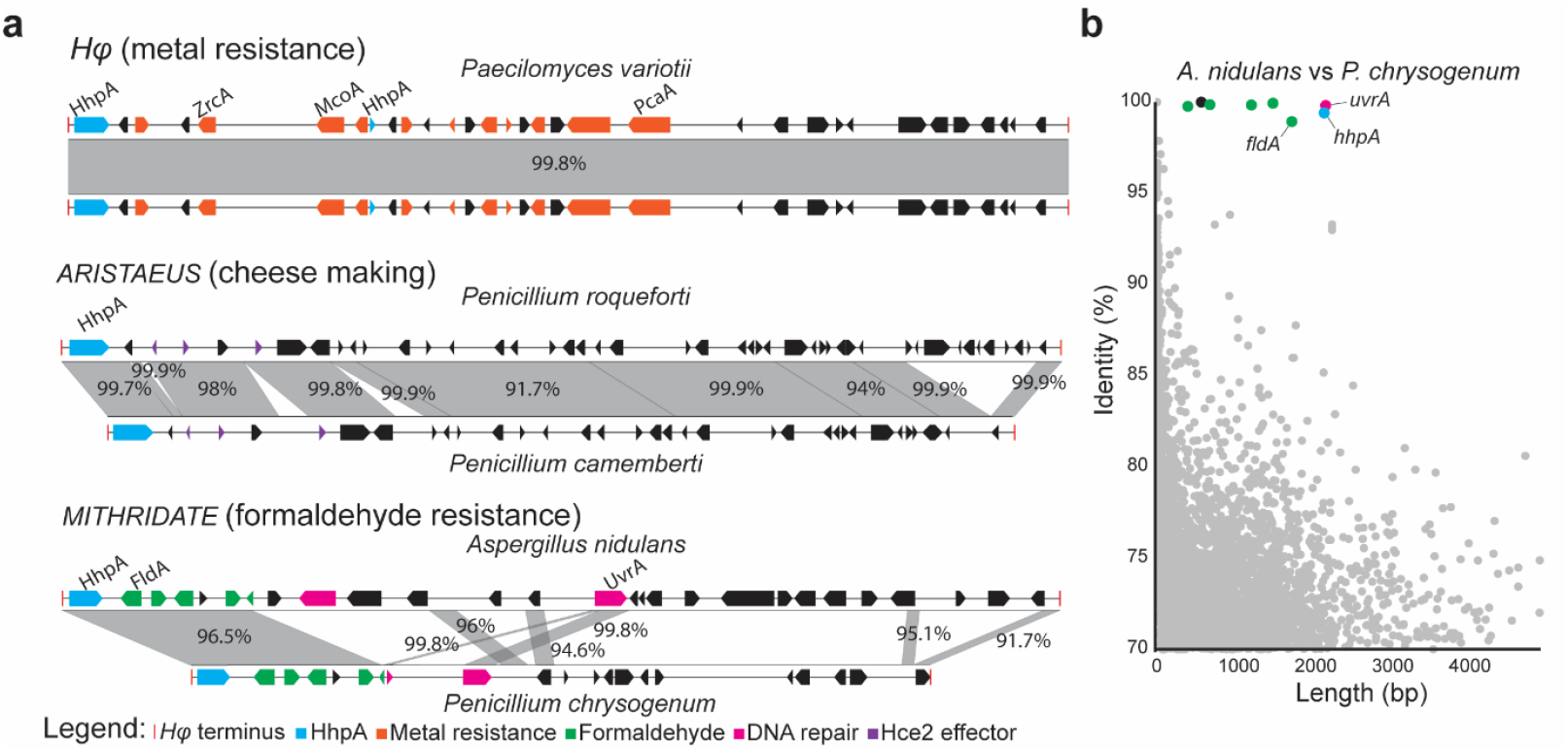
*HEPHAESTUS* family *Starships* have transferred diverse host-beneficial cargo between fungal species. **a**, Three examples of HGT events putatively driven by *Hφ-family* elements. HGT events involving metal resistance genes, including the original *Paecilomyces Starship*, were previously described. *ARISTAEUS* is a previously identified HGT event between cheese making fungi, but now annotated as an *Hφ* element. *MITHRIDATE* in *Aspergillus nidulans* strain SP-2605-48 contains an *Hφ-family Starship* featuring genes putatively involved in formaldehyde resistance and DNA repair. **b**, Many of the genes within *MITHRIDATE* show a strong HGT signal with a similar element in *P. chrysogenum* as assessed using BLAST all vs all comparison of the two genomes.

The first is an HGT previously identified between the cheese making fungi *Penicillium roqueforti* and *Penicillium camemberti* ^38^. Ropars et al. identified seven HGT events among the genomes of cheese making fungi in the genus *Penicillium*, which they designated as HGT 1 to 5, *Wallaby* and *CheesyTe*r. Recently, many of these have been annotated as *Starships* ^22^. One of these, HGT2, is a *Hφ-family Starship* which we name *ARISTAEUS* after the Greek god of cheese making. Consistent with other *Hφ*-family elements *ARISTAEUS* contains a *hhpA* homolog which is closely related to that of the original metal resistance *Hφ* of *P. variotii*. Furthermore, in both *P. roqueforti* and *P. camemberti* it has inserted into a TTAC(N7)A site (Fig. 3C). Comparison between the *ARISTEUS* elements of these two *Penicillium* species shows large regions close to 100% nucleotide conservation with regions of lower conservation, apparently the result of RIP. Ropars et al. have previously demonstrated that this high conservation is inconsistent with vertical inheritance from a common ancestor ^38^. Evidence suggests that at least some of the HGT regions within cheese fungi confer a fitness advantage both adapting to the cheese substrate and in out-competing other fungi ^38^. In particular, *Wallaby* contains a *paf* (*Penicillium* antifungal protein) gene identical to a *paf* in *P. rubens* which has been shown to be cytotoxic to fungi. *Wallaby* also encodes a second putatively antimicrobial protein which consists of Hce2 domain fused to a GH18 chitinase domain ^39,40^. Examination of the gene content of *ARISTAEUS* reveals a number of small Hce2 genes. These genes may, similar to genes in *Wallaby*, play a role in microbial competition ^40^.

A second example of a non-metal *Hφ*-family element is one putatively involved in formaldehyde resistance within the genome of *Aspergillus nidulans* strain SP-2605-48 ^41^. We name this element *MITHRIDATE* after the ancient poison antidote. Within *MITHRIDATE* at least five genes are putatively involved in formaldehyde resistance (Fig. 5A). In particular, *fldA* encodes a protein that is ~90% identical to *FldA* in *Paecilomyces formosus* NBRC 109023 (no. 5) that has been characterised as an S-hydroxymethylglutathione dehydrogenase involved in formaldehyde resistance ^42^.

This provides strong evidence that this cluster confers resistance to formaldehyde. This *Starship* is absent from other *A. nidulans* strains, including FGSC A4, allowing the target site to be defined as a typical *Hφ* TTAC site (Fig. 3C). A large section of this *Starship* shows HGT between *A. nidulans* and *P. chrysogenum* (Fig. 5B) with greater than 95% identity despite these species belonging to separate genera.

Two genes within *MITHRIDATE* are putatively involved in DNA repair, namely homologs of *rad4* and *uvrA*. UvrA shows similarity to bacterial UvrA homologs involved in DNA repair ^43^. This gene shows an unusual phylogenetic distribution with only a small number of fungal homologs and many closely related Gram-negative bacterial homologs. This is the second *Hφ*-family cargo gene to have been identified as being derived from a bacterial to fungal HGT, the other being a the *merA* and *merB* genes which detoxify mercury ^44^. A third gene encoding a uridine kinase in the *Hφ*-family element of *Aspergillus leproris* also appears to be a bacterial to fungal HGT. While we have no direct evidence for the involvement of *Starships* in bacterial to fungal HGT, *Starships* provide a mechanism by which such novel genetic information can be spread between fungal species.

The discovery of the *MITHRIDATE Starship* together with the annotation of *ARISTAEUS* as a *Starship* broadens the types of genes that the *Hφ-family* of *Starships* has horizontally transferred between species in the *Eurotiales*. We expect that as more elements are discovered and as more genes are characterised this role will continue to expand. The clustering of genes with a shared function (i.e., formaldehyde resistance in *MITHRIDATE* and putative Hce2 effectors in *ARISTAEUS*) mirrors the clustering of metal resistance genes in metal resistance *Hφ* elements. The grouping together of genes with a common function within the element is another parallel to bacterial ICEs ^8^ further supporting the hypothesis that the *Starships* and ICEs play analogous evolutionary roles in their respective kingdoms.

### Host beneficial *Starship* transposons are in tension with host genome defences that protect against mobile DNA

*Paecilomyces* has an active sexual cycle ^45^ in which repeat induced point mutation (RIP), a genome defence mechanism that mutates duplicated DNA (via C to T mutations), is known to be active ^27^. There are three possible situations in which *Starships* such as *Hφ* and *PEGASUS* would become vulnerable to RIP.

Firstly, entire *Starships* may be multicopy within genomes. Given that the position of *Hφ* is variable within the genome, it is possible for a strain to inherit two copies of *Hφ* following a cross. In such a situation we hypothesised that *Hφ* might become vulnerable to RIP mutation. Crosses were set up between wildtype strains to create a strain having two copies of *Hφ*, which was then crossed to a sibling with no copy. Rather than the expected 50:50 segregation of tolerant vs. sensitive, 29/30 progenies were more sensitive to zinc and 30/30 were more sensitive to cadmium ions compared to *Hφ+* strains (Fig 6A). This is despite many progenies inheriting one or both *Hφ* alleles (Fig 6B). DNA sequencing of a selection of these progenies revealed a pattern of transversion mutations (C/T or G/A) consistent with RIP (Fig. 6C). RIP frequently appeared to have acted in a strand specific manner (either predominately C -> T or G -> A mutations), such strand-specific patterns of RIP have been observed previously ^46^. The single progeny (#23) which retained full tolerance to zinc contained nonsynonymous mutations within the *pcaA* (conferring cadmium resistance) gene but not within the *zrcA* gene (conferring zinc resistance) consistent with the observed phenotype (data not shown).

**Fig. 6.**
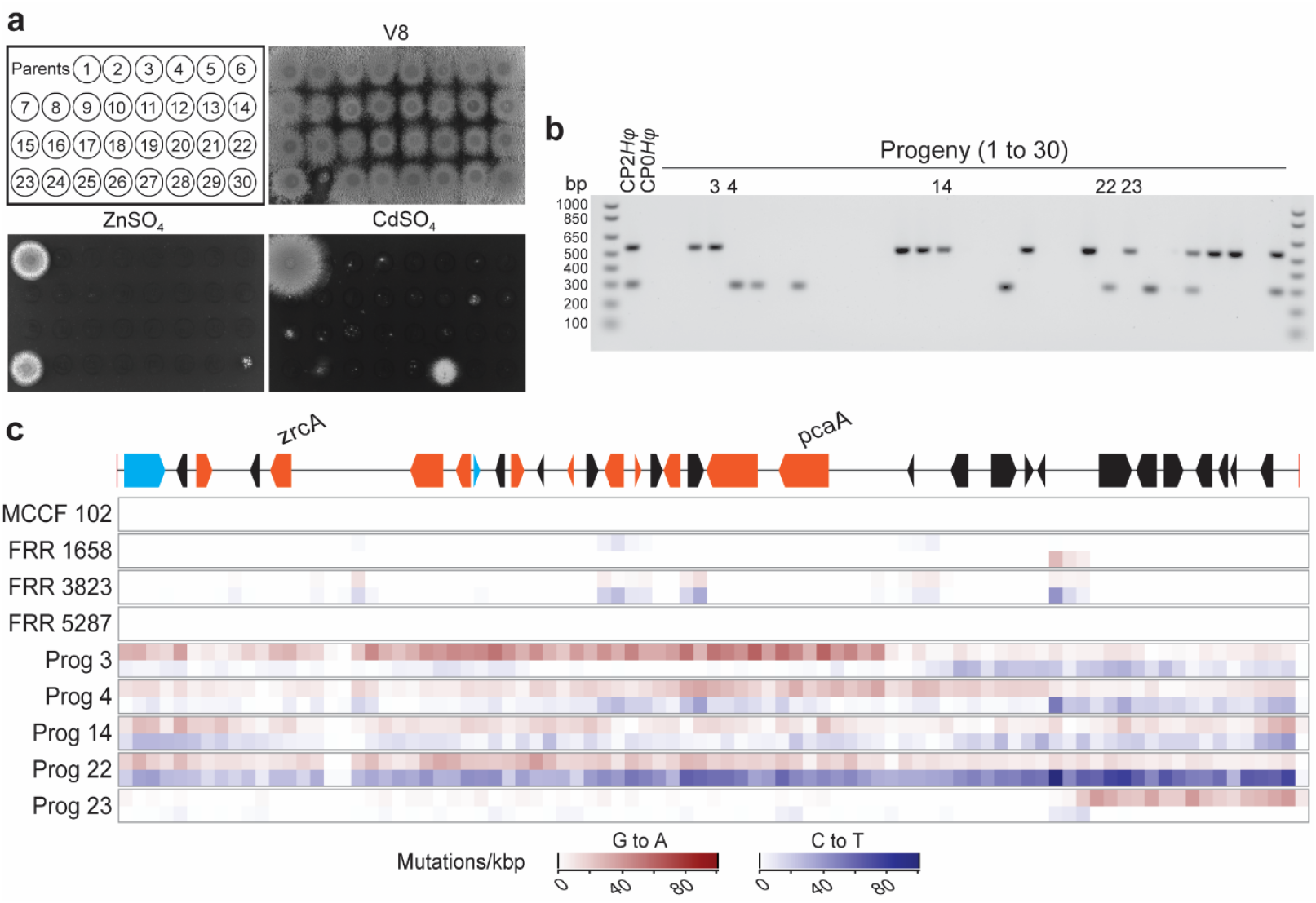
Mobility within the *Paecilomyces* genome leaves *HEPHAESTUS* vulnerable to destruction via RIP mutation. **a**, Metal ion sensitivity phenotypes of parents and the progeny when crossed. **b**, Despite showing sensitivity to zinc and cadmium, PCR analysis showed that 19 of the 30 progeny had inherited a copy of the *Hφ* region. **c**, Multiple sequence alignment of *Hφ* elements in wildtype strains of *Paecilomyces* or laboratory-generated progeny showing a high frequency of transversion mutations indicative of RIP. The positions of the *zrcA* and *pcaA* genes that confer resistance to zinc or cadmium, respectively, are indicated.

Secondly, *Starships* may contain internal duplications. Indeed, we found several internal repeats within *PEGASUS*. And we also found that *PEGASUS* was subjected to RIP during a cross (Fig. S5). This suggests that *PEGASUS* has entered this *Paecilomyces variotii* linage since the most recent instance of sexual reproduction. The frequency of sex among *Paecilomyces* strains in nature is not known and thus we cannot directly infer a maximum likely age of the element in this species, however, it has been shown that *P. variotii* has a recombining population structure, it readily mates under non-specific conditions (e.g. on potato dextrose agar), and the highly heat-resistant sexual spores have been suggested as a contributing factor to the prevalence of *P. variotii* in heat-treated foods ^45^. This all suggests that sex is likely to be a part of the biology of this species.

Thirdly, regions within *Starship* elements may show sequence similarity to the host genome. Wildtype *Hφ* sequences also show some evidence of RIP but this is localised to particular regions of the *Hφ* element (unlike RIP observed in the case of complete element duplication), perhaps due to similarity in these regions to the host DNA (Fig. 6C).

Host-beneficial *Starships* such as *Hφ* embody an inherent tension between their selfish properties – i.e., their mobility which is something which fungi typically employ defences against and their cooperative properties i.e., the host-beneficial genes they contain. Presumably both have been evolutionarily selected as advantageous to the *Starships* - in the first case by increasing the fitness of the *Starship* directly and in the second by increasing the evolutionary fitness of the whole organism. One might question the evolutionary benefit that mobility within the genome provides to a *Starship* which is carrying genuinely host-beneficial genes given that it leaves the *Starship* vulnerable to RIP (as we have demonstrated). Indeed, remnant *Starships* which have lost the ability to mobilise yet continue to persist in the host are common ^22^, these regions have essentially transitioned to being part of normal genomic landscape. We hypothesize that the resolution to this apparent paradox lies in considering horizontal transfer not as an oddity but as a fundamental feature in the lifecycle of host-beneficial *Starships*. That they mobilise primarily to allow a chance of landing in a different genome where the selective advantage of the novel genetic information they contain will drive their rapid spread through a new population. The types of phenotypes conferred within *Starships*: metal ion resistance ^21^, formaldehyde resistance (*MITHRIDATE*), pathogenicity ^22,37^ and potentially microbial competition (*ARISTAEUS*) show an emerging pattern towards resisting stresses which are heterogenous in the environment ^6,7^. This is in keeping with the idea of an inherently unstable evolutionary dynamic relying on the introduction of novel genic information, conferring strong selective advantages, into new populations for the survival of *Starship* elements. Such hypothesises provide another frontier for further exploration.

## Conclusion

Large DNA regions, dubbed ‘*Starships*’, have recently been hypothesized to act as host-beneficial eukaryotic transposons ^21,22^. Here we demonstrate the mobility of *Starship* elements and show that this movement is mediated by the ‘Captain’ tyrosine recombinase. Furthermore, the identification of a near-identical *Starship* in the genomes of two different fungal species is compelling evidence that these elements move not only within a genome but also between species. We thus provide firm experimental evidence for a role of *Starships* as an evolutionary force in fungi, analogous to ICE elements in bacteria.

We anticipate that the true impact of *Starships* in the mobilisation of host-beneficial DNA within and between species remains hidden. As Arkhipova and Yushenova ^19^ point out, ‘there may be only a short window of time for bona fide HGT mediated by TEs to be identified.’ The examples of HGT provided here likely represent relatively recent events as they can be detected straightforwardly by unexpectedly high DNA sequence conservation. More ancient events, in which significant divergence have occurred, could not be detected by such an approach and will likely require more careful analysis of gene phylogenies to infer HGT. As noted previously, in many such cases, the TIRs or Captain recombinases many have been lost, and the involvement of *Starship* in those events would be more difficult to recognize ^21,22^. Thus, we have only scratched the surface of the role of *Starship*-mediated HGT in fungi, i.e. by examining the most obvious cases in a single *Starship* family - *Hφ*.

Bioinformatic exploration of genome sequences clearly indicates that *Starships* are a pervasive phenomenon in at least the Pezizomycotina fungi ^22^. Already implicated in chemical resistance, pathogenicity, and fungal domestication, the study of *Starships* represents an exciting avenue towards increasing our understanding of how fungi rapidly evolve numerous traits that impact upon human activities

## Materials and Methods

### Strains

*Escherichia coli* strain DH5α was used for molecular cloning. *Agrobacterium tumefaciens* strain EHA105 was used for *Agrobacterium* mediated transformation. *Saccharomyces cerevisiae* strain BY4742 was used for yeast assembly of DNA constructs. *Paecilomyces* strains used and generated in this study are listed in Table S5.

### DNA cloning

#### PLAUB36 (pPZP-201BK adapted for yeast cloning)

The DG1246+DG1247 PCR fragment of pYES2, containing the *URA3* gene conferring uracil prototrophy and the 2-micron origin allowing plasmid replication in yeast, was cloned into the ScaI site of pPZP-201BK ^47^ using homologous recombination in yeast. This allows yeast cloning into an *Agrobacterium* vector. Yeast cloning was done by introducing the linear DNA fragments into *S. cerevisiae* cells using a LiAc/PEG method ^48^ with selection on SC media without uracil. The linear fragments were combined into a circular plasmid by the native *S. cerevisiae* homologous recombination machinery and rescued into *E. coli* using the Zymoprep Yeast Plasmid Miniprep II kit (Zymo Research).

#### PLAUB63 and 64 (Constructs made to “double flank” the native *Hφ* copy)

PLAU64 - A fragment 5’
s to *Hφ* was amplified with primer pair AUB314+AUB315. The G418 marker was amplified from a previously generated construct MT680156 ^49^ with AUB316+AUB317, The actin promoter of PMAI6 ^50^ was amplified with primers AUB318+AUB319. The 5′ edge of *Hφ* was amplified with AUB320 and AUB321. These four fragments were combined into plasmid PLAUB36 linearised with EcoRI and HindIII as described above.

PLAU63 - The 3’
s edge of *Hφ* was amplified with AUB322+AUB323. The hygromycin phosphotransferase cds and terminator was amplified from PMAI6 ^51^ using primers AUB324+AUB325. The nours marker was amplified from a previously generated construct MK431404 ^52^ using primers AUB326+AUB327. A genomic fragment 3′ to *Hφ* was amplified with primer pair AUB328+AUB329. These 4 fragments were combined into plasmid PLAUB36 linearised with EcoRI and HindIII as described above.

Primer pair AUB412+AUB413 were used to verified integration of the G418 construct and AUB414 and AUB415 for the nourseothricin resistance construct.

#### PLAUB28 (*Hφ*mini)

The promoter driving expression of the hygromycin phosphotransferase (*hph*) gene was amplified with primers AUB244 and AUB251 off PMAI6 ^51^. The 5’
s side of *Hφ* including the TIR and HhpA was amplified with AUB215 and AUB216. The *zrcA* gene conferring zinc resistance with *Hφ* was amplified with AUB253 and AUB254. The 3′ side of *Hφ* was amplified with primers AUB219+AUB220. The *hygR* open reading frame and terminator was amplified from PMAI6 ^51^ using primers AUB252 and AUB243. These five fragments were combined into plasmid PLAUB36 linearised with EcoRI and HindIII as described above.

#### PLAUB71 (actin promoter: HhpA) and PLAUB80-PLAUB83 (actin promoter: HhpA point mutations)

The HhpA coding region was amplified with primer pair AUB424 + AUB425 from plasmid PLAUB28 and cloned into the BglII restriction site of PLAU53 ^51^ using the NEBuilder Assembly Kit which results in HhpA expression under control of the constitutive *Leptosphaeria biglobosa* actin promoter and G418 resistance to allow selection of strains transformed with the T-DNA.

Three plasmids identical to PLAUB71 except for point mutations within *hhpA* were created. Mutated versions of *Hφ* were generated in two halves with primer pairs AUB424+AUB512 and AUB425+AUB513 (Y381A); AUB424+AUB515 and AUB425+AUB514 (Y381H);and AUB424 + AUB496 and AUB425+AUB497 (Y416A) and combined into the BglII site of plasmid PLAU53 ^51^ using the NEBuilder Assembly Kit.

#### PLAUB69 (HhpA GFP tagging construct)

A fragment of the 3’
s end of the *hhpA* coding region was amplified using AUB416 and AUB417, GFP coding DNA was amplified from plasmid PLAU17 ^51^ with primer pair AUB418 + AUB419, the hygromycin resistance cassette of plasmid PMAI6 ^51^ was amplified with primer pair AUB420 + AUB421 and a section of genomic DNA 3′ of *hhpA* was amplified with primer pair AUB422 + AUB423. The four regions were combined into plasmid PLAUB36 linearised with EcoRI and HindII using homologous recombination in yeast.

#### PAUB72 (H2B-mScarlet)

A DNA fragment encoding a histone H2B-Scarlet ^53^ fusion protein was chemically synthesised by IDT and cloned into the BglII site of plasmid PLAU53 ^51^ using the NEBuilder Assembly Kit. The synthesised sequence is given in supplemental Fig. S6. Design of this construct is based on a previous histone H2B-CFP construct ^27^.

### Sexual crosses

Crossing was conducted as described previously ^27,45^. Briefly strains of opposite mating type were streaked approximately 20 mm apart on 90 mm potato dextrose agar plates. Strains were incubated for up to six weeks at 30°C until sexual spores could be observed. Frequently these form at the site of initial inoculation rather than at the junction of the two strains. Sexual spores were harvested into water and then incubated at 80°C for 10 mins, which is sufficient to kill asexual spores of *P. variotii* ^45^. The heat-treated spores were plated onto fresh PDA plates and incubated for 24-48 h until colonies emerged which were further purified by single-spore technique. Progeny were screened for mating type by duplex PCR with primers MAI0448, MAI0449, MAI0450 and MAI0451 ^27^, and *Hφ* using a duplex PCR targeting polymorphic host DNA adjacent to the insertion site with primers AUB542, AUB544, AUB546 and AUB547 (298 bp FRR 3238 insertion, 517 bp CBS 144490 insertion).

### *PEGASUS* segregation analysis

Twenty-two progeny from a cross between CBS 144490 x CBS 101075 were previously obtained and characterized for segregation of *Hφ* and the mating type locus ^21^. The progeny were cultured from - 80°C glycerol stocks, genomic DNA extracted, and fragments corresponding to different contigs amplified by PCR using TaKaRa Ex Taq DNA polymerase with primers MAI830 to MAI870 (Table S6). Products were resolved on 1% agarose gels and scored for the presence/absence of bands.

To detect RIP mutation within *PEGASUS* in these progenies, a part of *PEGASUS* (4.3 kb) was amplified from wild type and two progeny (numbers 3 and 10) by PCR with MAI0857 and MAI0858. The amplicons were sequenced with these and several internal primers. The sequences were deposited to GenBank as ON622788 (wildtype), ON774807 (Progeny 3) and ON774808 (Progeny 10).

### Taxonomic determination of strains of *Paecilomyces* with sequenced genomes

Sequenced *Paecilomyces* genomes were identified through BLASTn searches of the NCBI whole-genome shotgun database. Misidentification of sequenced genomes is a recognised issue ^54^ and thus we taxonomically categorised these genomes to the species level using a fragment of the calmodulin gene. The calmodulin gene sequence was extracted from the assemblies via BLASTn and compared to taxonomically certain strains from ^35^. Sequences were aligned using Clustal Omega ^55^ and a phylogeny was inferred using a Bayesian approach implemented in MrBayes using the HKY85 substitution model ^36^. We followed the taxonomic scheme proposed by Samson et al. ^35^ except that we have treated *P. lecythidis* and *P. maximus* ^56^ as separate from *P. formosus*, because molecular evidence supports the separation of these species even in the absence of clear morphological support ^35^. To minimise confusion, we have renamed strains to reflect their correct taxonomy (Table S7).

### Fungal transformation

*Paecilomyces* was transformed using *Agrobacterium* as described previously ^27^. Briefly, plasmids (~200 ng in 2 µl) were electroporated into *Agrobacterium* cells (50 µl ~10^6^) that had first been washed 3 times with cold 10% glycerol. Electroporation was performed using a BioRad Gene Pulser II (2 kV, 25 µF, 500 ohms). Transformed cells were recovered in SOC medium for ~2-3 h then plated onto LB media supplemented with kanamycin. After colonies had formed *Agrobacterium* was scrapped directly from the transformation plate and resuspended in SOC (~0.5 OD). 100 µl of *Agrobacterium* suspension was mixed with approximately 1 × 10^6^ fungal spores on a 15 cm petri dish containing 25 ml of *Agrobacterium* induction media ^57^. After three days incubation the plates were overlayed with 25 ml of cleared V8 media supplemented with approximated selective agent (100 µg/ml hygromycin, 100 µg/ml G418, 200 µg/ml nourseothricin or 5 mM ZnSO_4_; the later allowing the use of the *zrcA* gene as a novel selectable marker) and 100 µg/ml cefotaxime to inhibit *Agrobacterium*.

### Transposition assay

Strains in which *Hφ* elements were “sandwiched” between a promoter and the hygR coding region were generated either with PLAUB63+PLAUB64 (to flank the native copy) or PLAUB28 (carrying the *miniHφ* allele). Excision of *Hφ* results in the promoter driving hygR expression leading to resistance to hygromycin. A single spore was placed at the centre of a 9 cm V8 agar petri-dish and incubated at 30°C for 3 weeks. The spores were harvested and approximately 10^8^ spores were replated on a fresh plate supplemented with 100 µg/ml hygromycin. Hygromycin resistant colonies were counted after 48 h incubation.

### Genome assembly of *Paecilomyces* strains

Illumina sequencing reads were previously generated for three *P. variotii* strains containing *Hφ* (FRR 1658,FRR 3823 and FRR 5287) ^21^. In order to examine the *Hφ* elements within these strains for signs of RIP mutation we assembled the genomes using Velvet ^58^ with a *k*-mer value of 89. In the three strains that contain a *Hφ* element the region was resolved into a single scaffold. The assemblies have been deposited in GenBank (Table S7).

### Presence of *Hφ* and *PEGASUS* in *Paecilomyces variotii*

The presence of the two *Starship* elements in previously sequences strains was assessed using BLAST searches. In four strains FRR 1658, FRR 2889, FRR 3823 and CBS 101075 this presence/absence of *Hφ* was confirmed by read mapping to the CBS 144490 assembly using bowtie2 ^59^. CBS 101075 was represented by short read sequencing generated from a *pvpP* mutant of CBS 101075 (manuscript in preparation).

### Metal stress testing

Tolerance to two metal ions Cd^2+^ and Zn^2+^ was determined according to the fungus’s ability to grow on 90 µM CdSO_4_ or 10 mM ZnSO_4_. These concentrations were chosen because *P. variotii* strains lacking a functional copy of the *pcaA* or *zrcA* genes, respectively, are unable to grow at these concentrations ^21^. Spores were plated onto cleared V8 media supplemented with the metal salts and incubated for 24 h at 30°C after which time growth was scored.

### Genomic sequencing

Genomic DNA was extracted using a CTAB based method as described previously ^60^. DNA was sequenced using an Illumina NovaSeq 6000 system with 150 bp paired end reads at the Victorian Clinical Genetics Services (Melbourne). Illumina reads were deposited in the SRA database (Table S8)

### AlphaFold protein structure predictions

Protein structures were for HhpA and KIRC were predicted using AlphaFold. The structures were aligned to each other or to known YR crystal structures using the pairwise structural alignment tool available through the RCSB PDB web portal (https://www.rcsb.org/alignment) using the default jFATCAT algorithm ^61^.

### Phylogenetic analyses of Captain proteins

Sequences with homology to *HhpA* were identified from Genbank with BLASTn The *Hφ-*family was delimited based on those sequences that formed a monophyletic group with *HhpA* and were largely restricted to the Eurotiales, though notable exceptions exist (Figure S1). Alignments were generated using MAFFT with default parameters. The phylogeny was constructed using IQ-TREE version 2.0.3 with automatic model selection and 1000 ultrafast bootstraps. The JTT+R5 model was selected.

A *hhpA* gene tree for *Hφ-*family starships with defined insertion sites was generated in MEGAX (using the K2+G+I model) from a nucleotide alignment manually edited to remove poorly aligned regions. Boot strap supports were generated from 1000 replicates.

### *Starship* gene content analysis

To determine the putative functions of *Hφ-*family cargo genes, eggNOG-mapper version 2.1.5 was run with default parameters. As no training set was given, the gene annotations were largely fragmented at introns. To address this, only the number of unique pfam domains was evaluated for each element, rather than gene content.

## Acknowledgements

We thank Dr Chunhong Chen for assistance running AlphaFold software. AU was supported by a CSIRO CERC postdoctoral fellowship. AV was supported by FORMAS 2019-01227 and the Swedish research council 2021-04290.

---

**Fig. S1.**
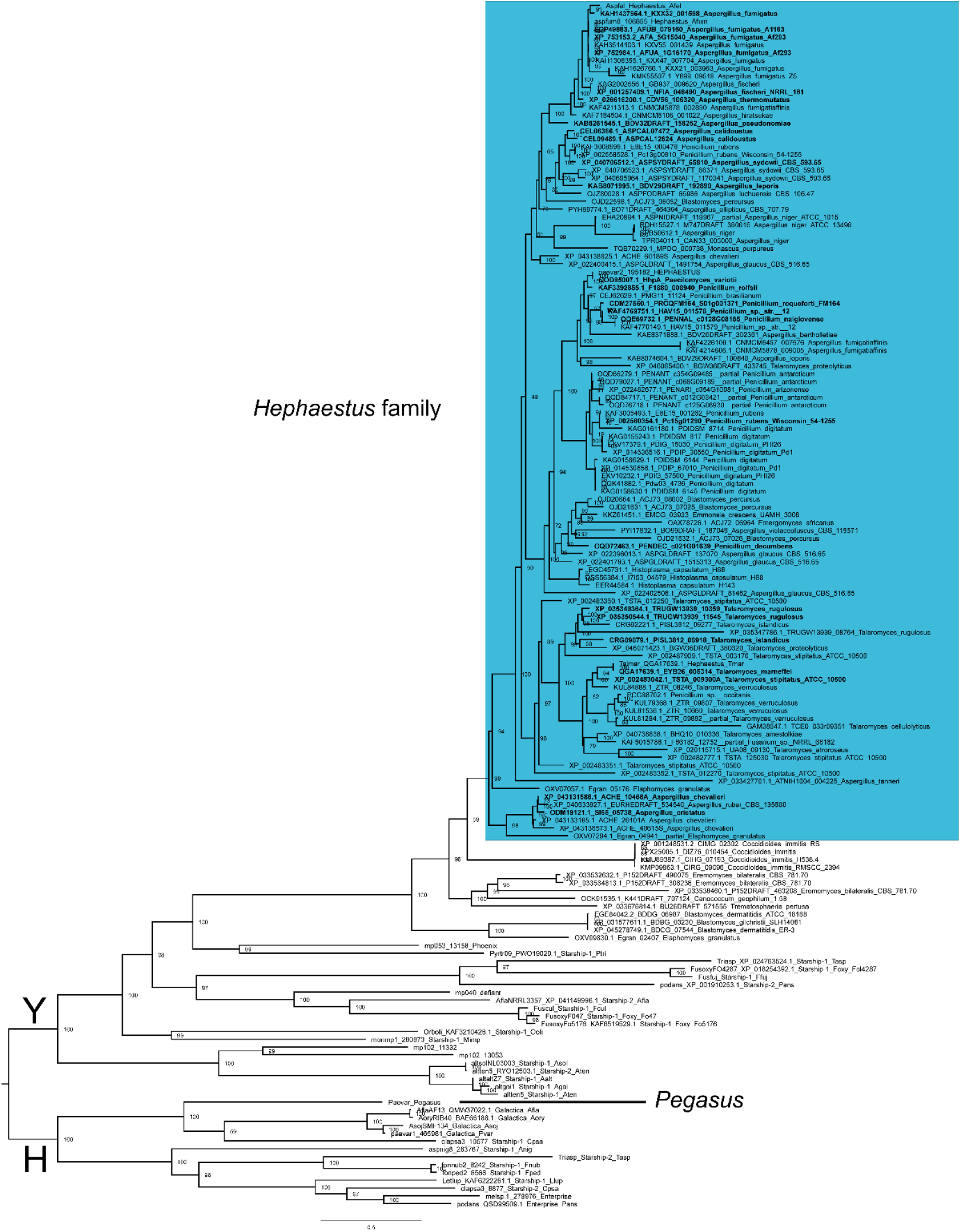
Phylogeny of *Starship* Captain proteins. Tree generated in IQ tree based on amino acid sequences. There is a split in family members depending on the presence of either tyrosine (Y) or histidine (H) at a conserved active site. Captains of elements that are fully assembled and clearly delimited (based on the presence of TIRs) are in bold. Nodes labels indicate support based on 1000 ultrafast bootstraps.

**Fig. S2.**
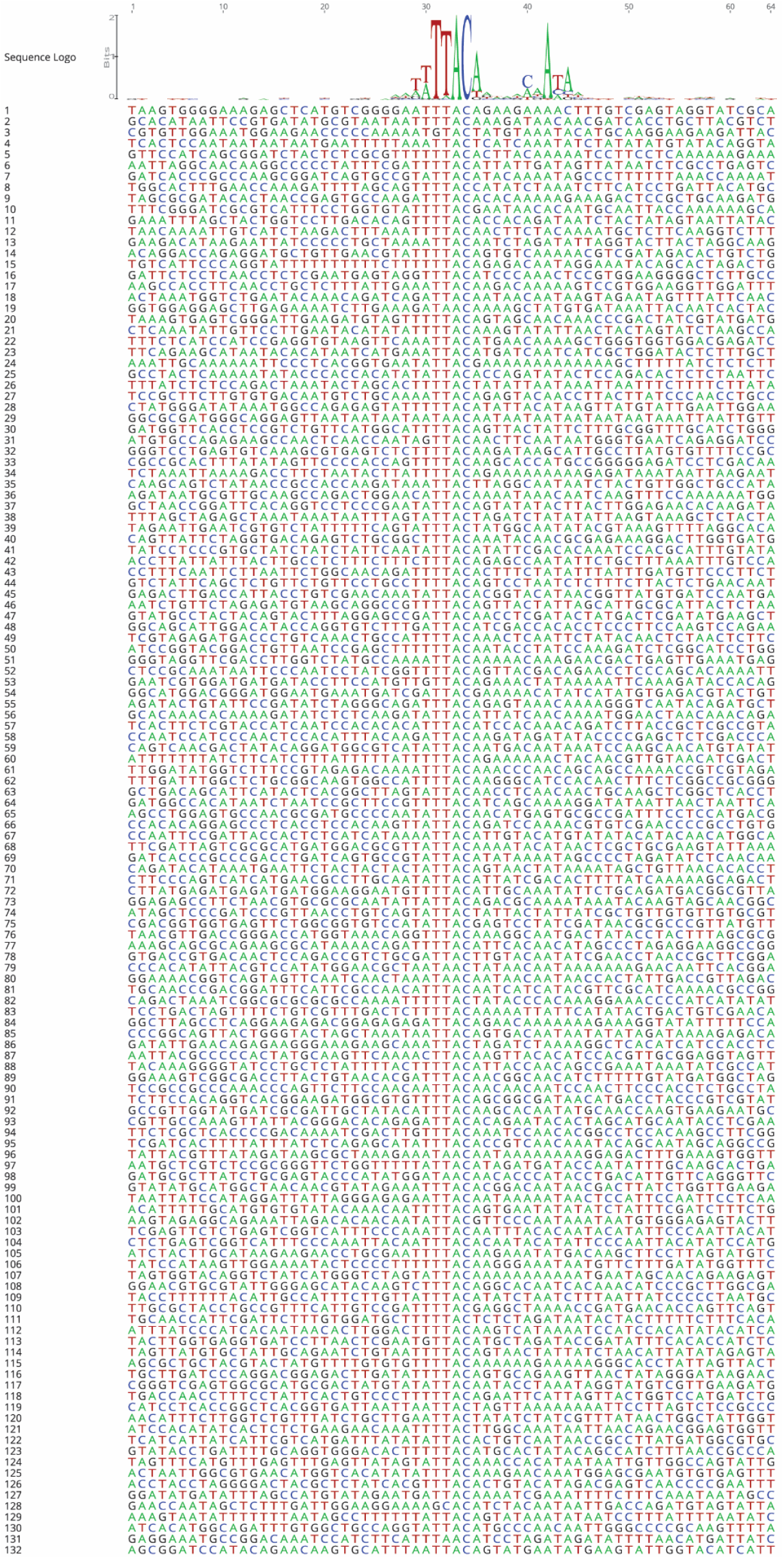
Consensus and alignment of 132 *miniHφ* insertion sites.

**Fig. S3.**
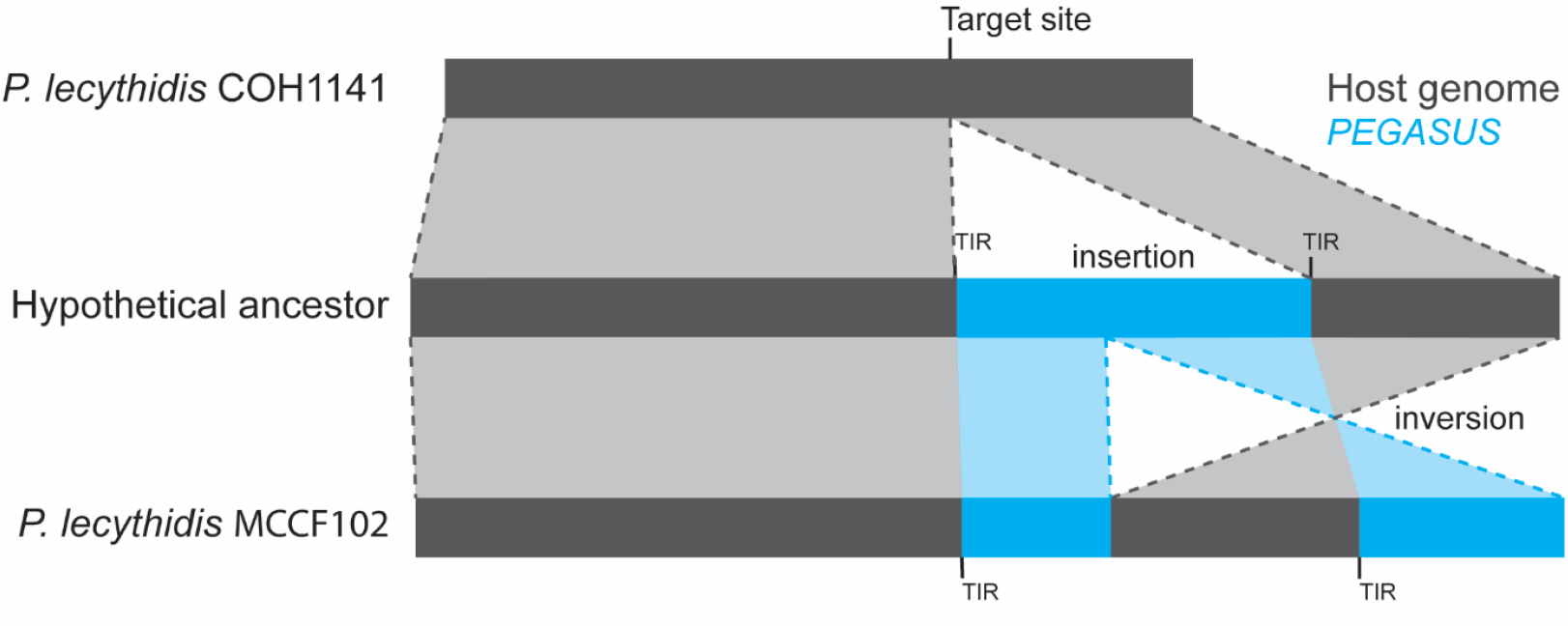
Inversion mutation in *PEGASUS* of *P. lecythidis* strain MCCF 102.

**Fig. S4.**
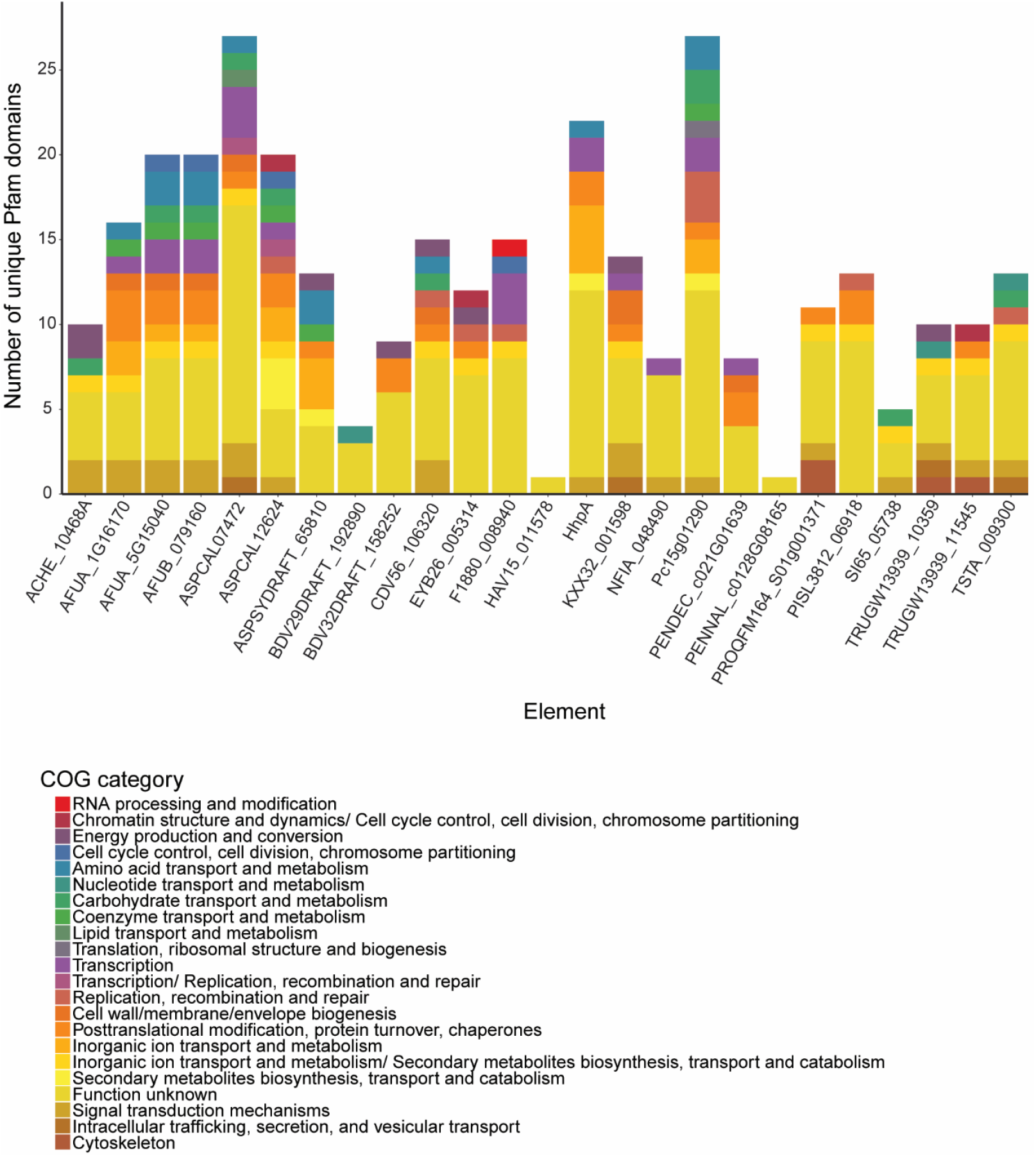
The gene content of 25 *Hφ-family* elements annotated by COG category.

**Fig. S5.**
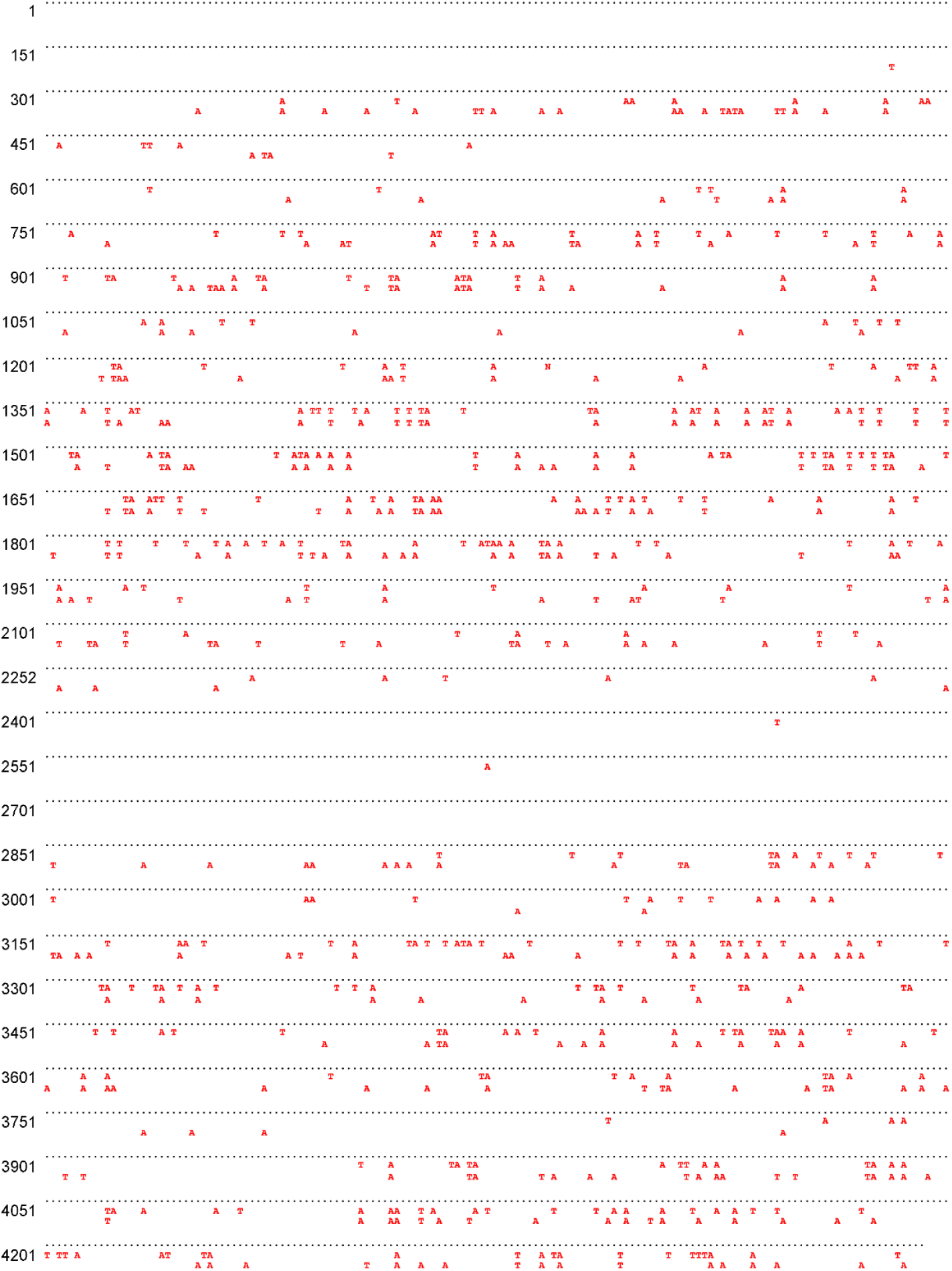
Illustration of the RIP mutations in a 4.3 kb fragment of *PEGASUS*. The wild type sequence is at the top, portrayed as dots, with underneath the sequences from progeny 3 and 10 whereby any C-T or G-A change is given in red font (GenBank accessions ON622788, ON774807 and ON774808). The numbers to the left indicate the base pair numbers. In total 381 and 360 C to T or G to A differences compared to the wild type were observed, i.e. RIP mutation changed 8.8% and 8.3% of the sequence, respectively.

**Fig. S6.**
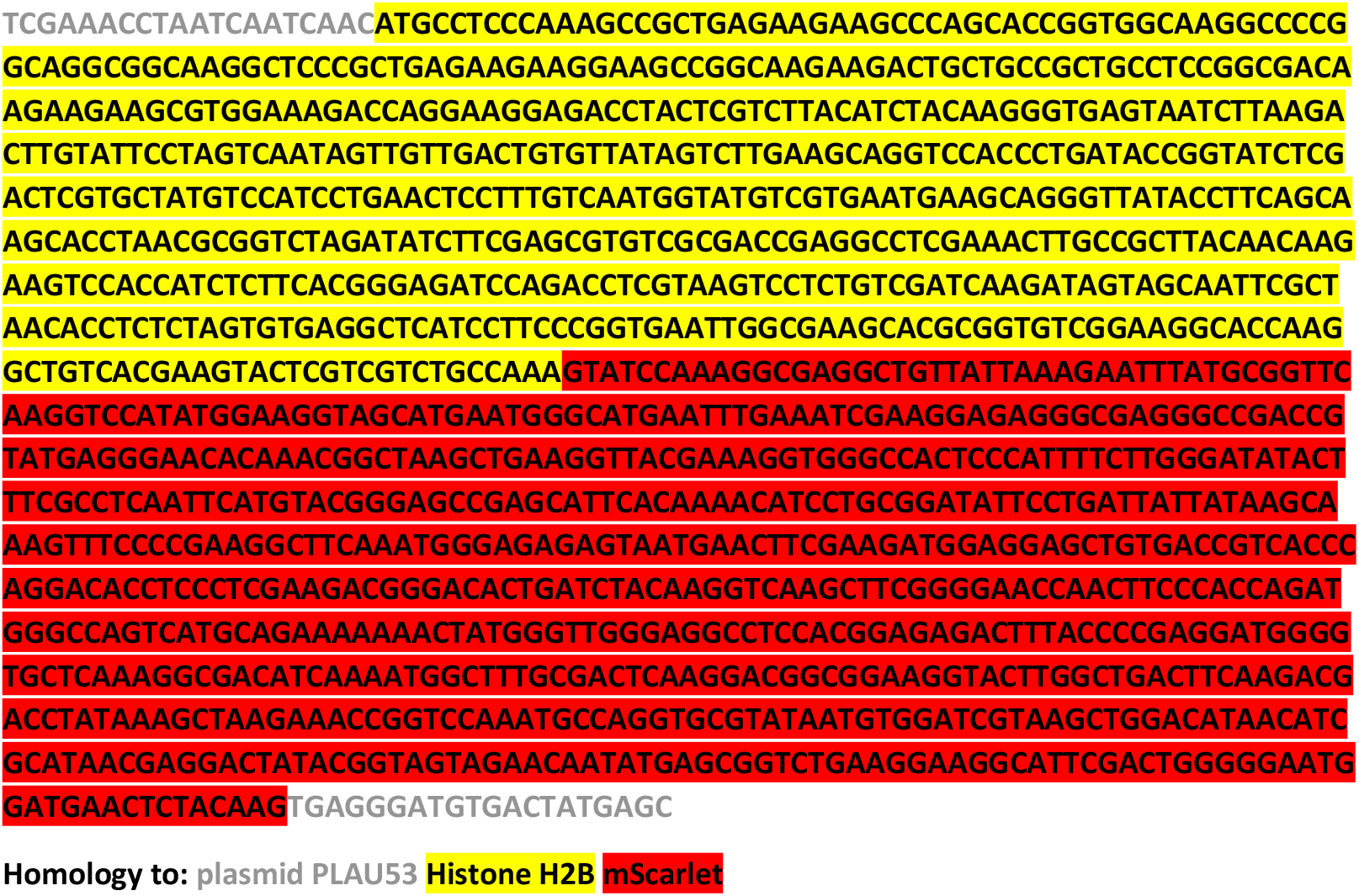
Sequence of the synthesised histone H2B-mScarlet fragment.

**Table S1.**
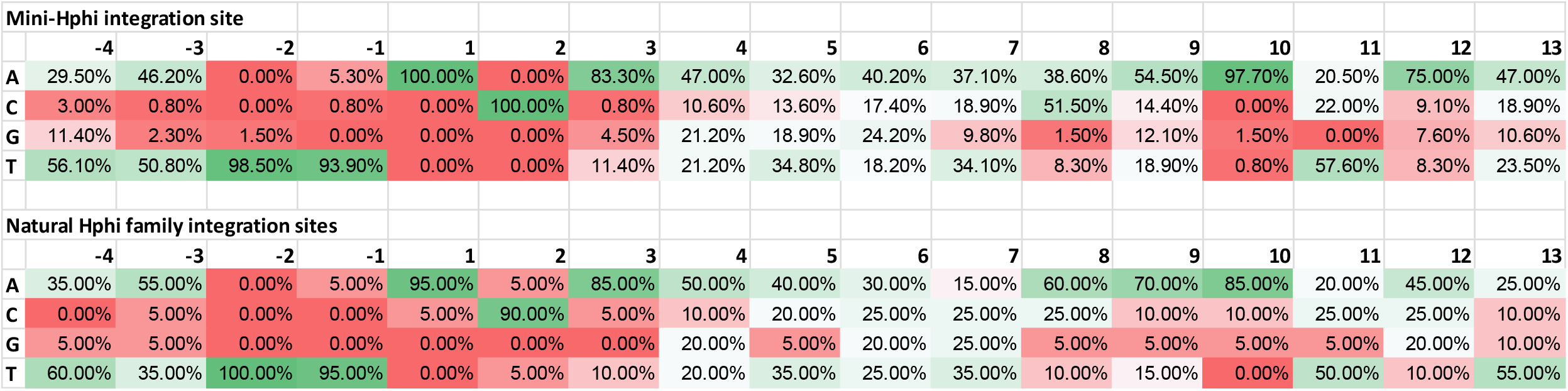
Nucleotide frequencies surrounding *Hφ* insertion sites.

**Table S2.** Newly identified *Hφ* relatives define based on comparison to a corresponding empty site.

| **Contig containing *Hϕ* element** | **Empty reference contig** | **Citations** |
| --- | --- | --- |
| *Aspergillus nidulans* SP-2605-48<br>JAAFYL010000042.1 | *Aspergillus nidulans* FGSC A4<br>BN001303.1 | 41,62 |
| *Aspergillus fumigatus* D-2-23L-5<br>JAIBQD010000015.1 | *Aspergillus fumigatus* Af293<br>CM000176.1 | 63,64 |
| *Aspergillus fumigatus* F-1-33L-2<br>JAIBXQ010000101.1 | *Aspergillus fumigatus* D-2-32L-1<br>JAIBZD010000134.1 | 65 |
| *Aspergillus lentulus* IFM 47457<br>BLKD010000031.1 | *Aspergillus lentulus* IFM 54703<br>BCLY01000009.1 | 66 |
| *Aspergillus fumigatus* Af293<br>AAHF01000003.1 | *Aspergillus fumigatus* A1163<br>DS499597.1 | 64,67 |
| *Talaromyces marneffei* 11CN-20-091<br>CP045656.1 | *Talaromyces marneffei* TM4<br>CP015873.1 | 68 |
| *Penicillium camemberti* FM013<br>JACXYV010000008.1 | *Penicillium camemberti* IIF8SW-F3<br>JACSOO010000009.1 | 69,70 |
| *Penicillium rubens* IMI 15378<br>CACPRF010000008.1 | *Penicillium chrysogenum* NCPC10086<br>KI963894.1 | 71,72 |
| *Penicillium roqueforti* FM164<br>HG792015.1 | *Penicillium roqueforti* JCM 22842<br>BCID01000001.1 | 40 |

**Table S3.**
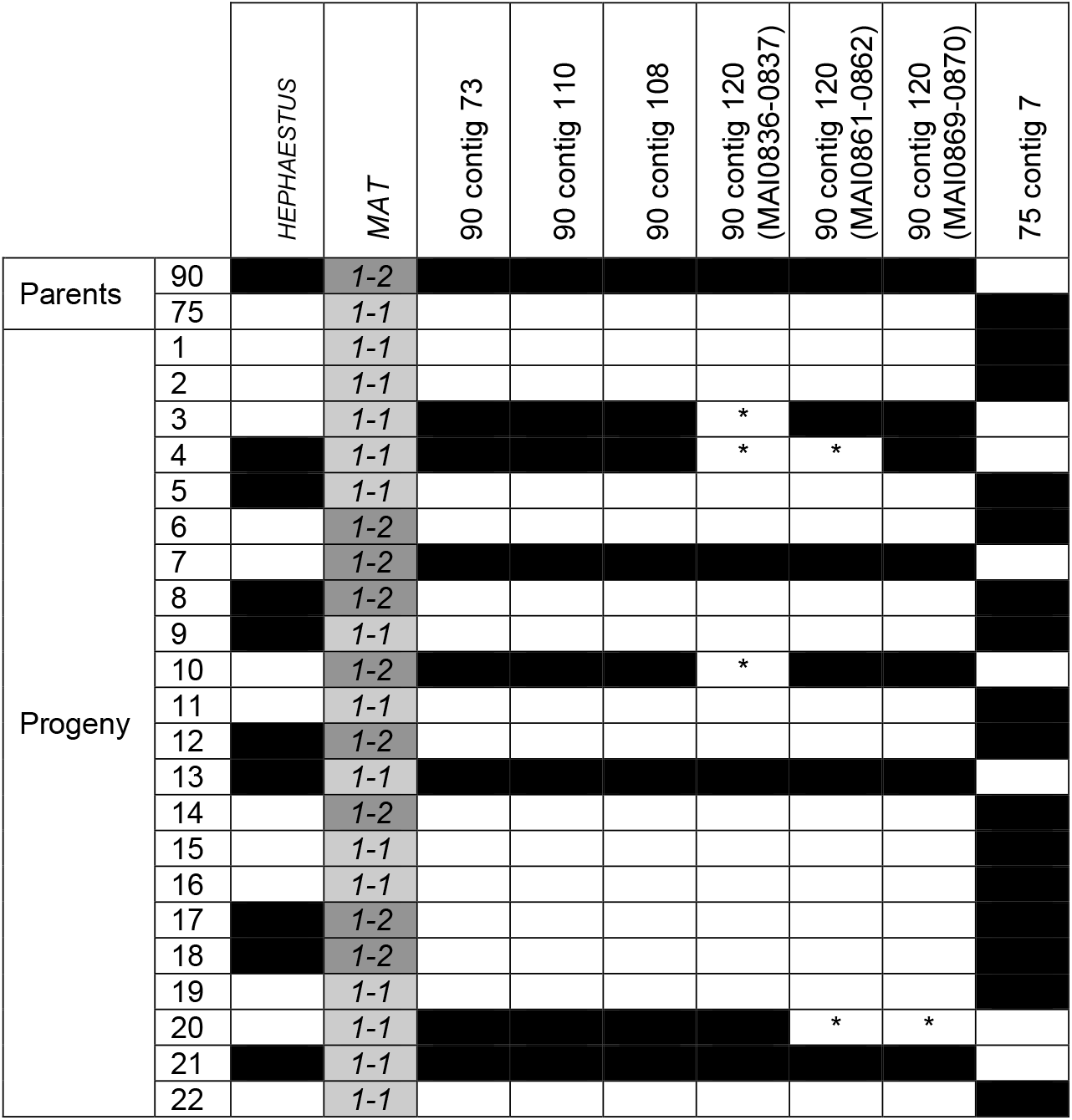
Segregation of molecular markers in 22 progeny (1-22) of a cross between CBS 144490 (90) and CBS 101075 (75). Black fill indicates amplification; the *MAT* locus segregates as two different idiomorphs, either *MAT1-1* or *MAT1-2*. * indicates no amplification, despite other PCRs on the same contig working (likely due to RIP mutation).

**Table S4.**
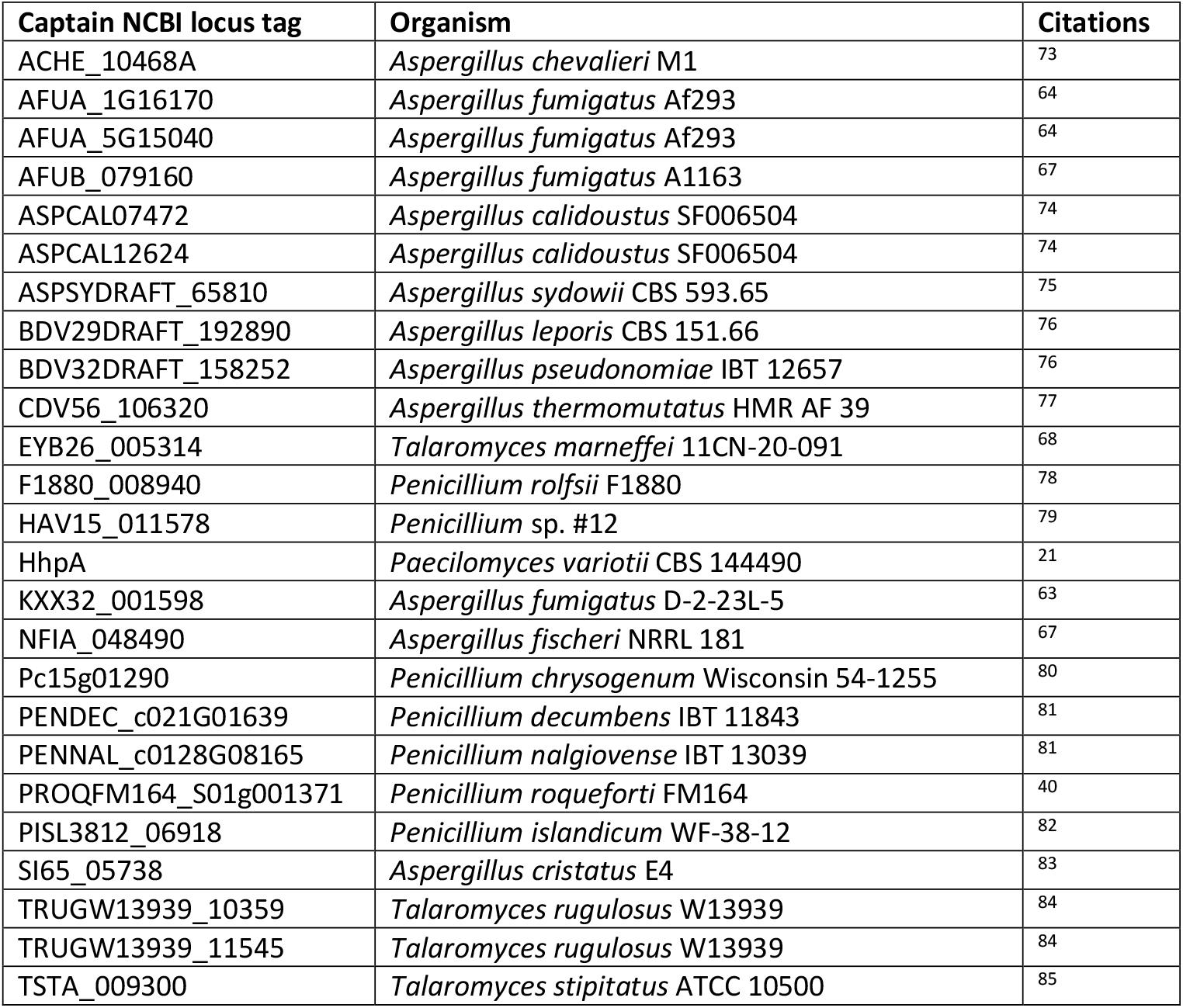
25 *Hφ-family* elements identified through a gene cargo neutral approach.

**Table S5.** Strains used in this study.

| Strain | Description | Source |
| --- | --- | --- |
| CBS 144490 (*Hφ*+) | WT strain (*Hφ*+) MAT1-1 | 27 |
| CBS 101075 (*Hφ*-) | WT strain (*Hφ* -) MAT1-2 | 27 |
| FRR 2889 | WT strain (*Hφ* -) MAT1-2 | 21 |
| FRR 3823 | WT strain (*Hφ*+) MAT1-1 | 21 |
| CBS 144490 x CBS 101075 progeny | 22 progeny from a cross between the two parents | 21 |
| CP2*Hφ* | Strain derived through sexual crosses. FRR 2889 was crossed to CBS 144490. An *Hφ*+ progeny of this cross was in term crossed to FRR 3823. CP2*Hφ* was a progeny of this cross containing both the CBS 144490 and FRR 3823 derived copies of *Hφ* and the MAT1-1 mating type. | This study |
| CP0*Hφ* | Strain derived through sexual crosses. FRR 2889 was crossed to CBS 144490. An *Hφ*+ progeny of this cross was in term crossed to FRR 3823. CP2*Hφ* was a progeny of this cross containing neither the CBS 144490 nor FRR 3823 derived copies of *Hφ* and has the MAT1-2 mating type. | This study |
| CP2*Hφ* x CP0*Hφ* progeny | 32 progeny of a cross between CP2*Hφ* and CP0*Hφ*. | This study |
| CBS 101075 *miniHφ* | CBS 101075 transformed with plasmid PLAUB28 | This study |
| CBS 101075 *miniHφ*-frameshift | CBS 101075 transformed with plasmid PLAUB56 | This study |
| CBS 101075 *miniHφ*-frameshift + HhpA<sup>WT</sup> | CBS 101075 *miniHφ* transformed with plasmid PLAUB71 | This study |
| CBS 101075 *miniHφ*-frameshift + HhpA<sup>Y381A</sup> | CBS 101075 *miniHφ* transformed with plasmid PLAUB81 | This study |
| CBS 101075 *miniHφ*-frameshift + HhpA<sup>Y381H</sup> | CBS 101075 *miniHφ* transformed with plasmid PLAUB82 | This study |
| CBS 101075 *miniHφ*-frameshift + HhpA<sup>Y416A</sup> | CBS 101075 *miniHφ* transformed with plasmid PLAUB83 | This study |
| CBS 144490 double flanked | CBS 144490 transformed with plasmids PLAUB63 and PLAUB64 | This study |

**Table S6:**
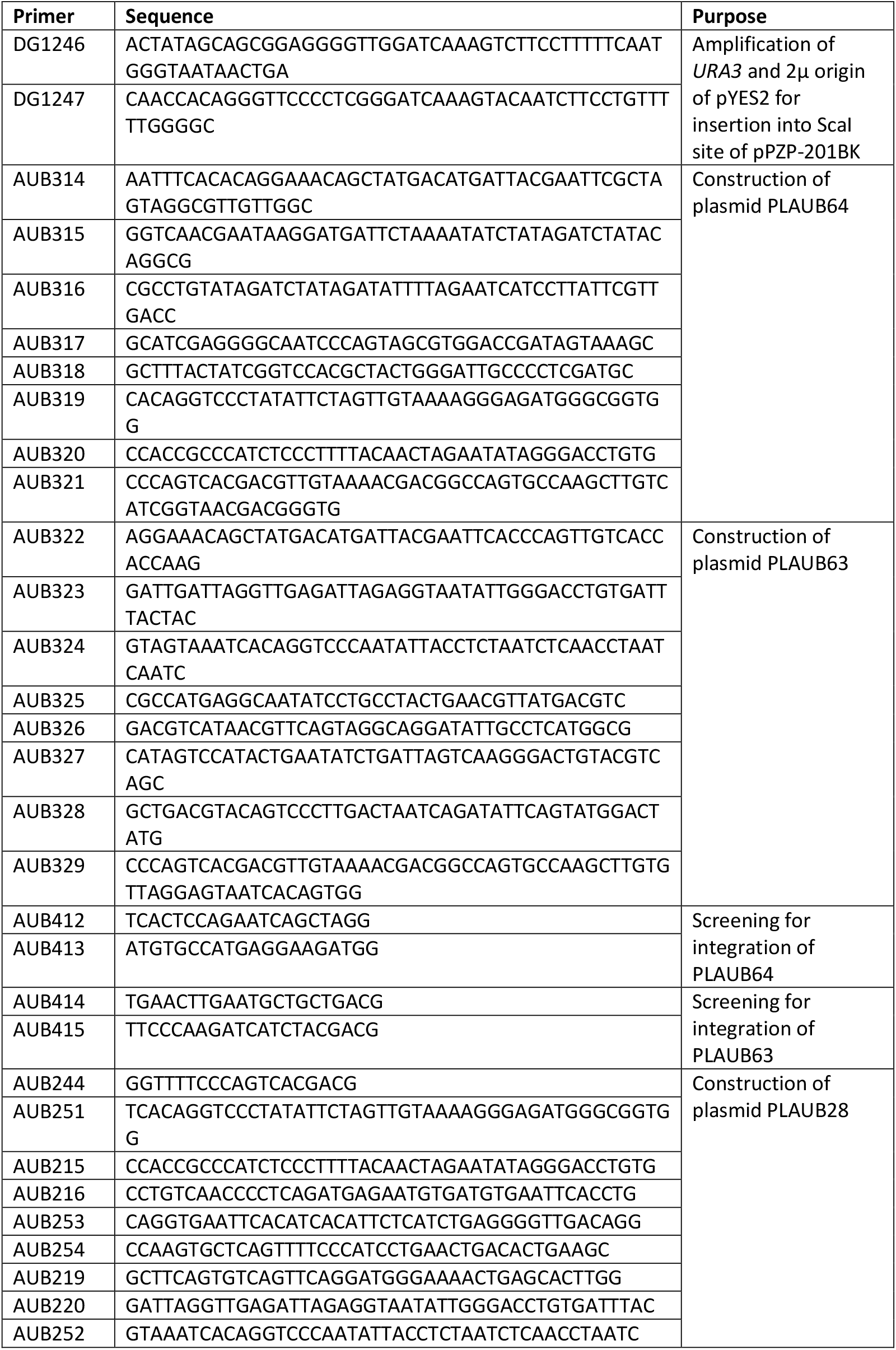
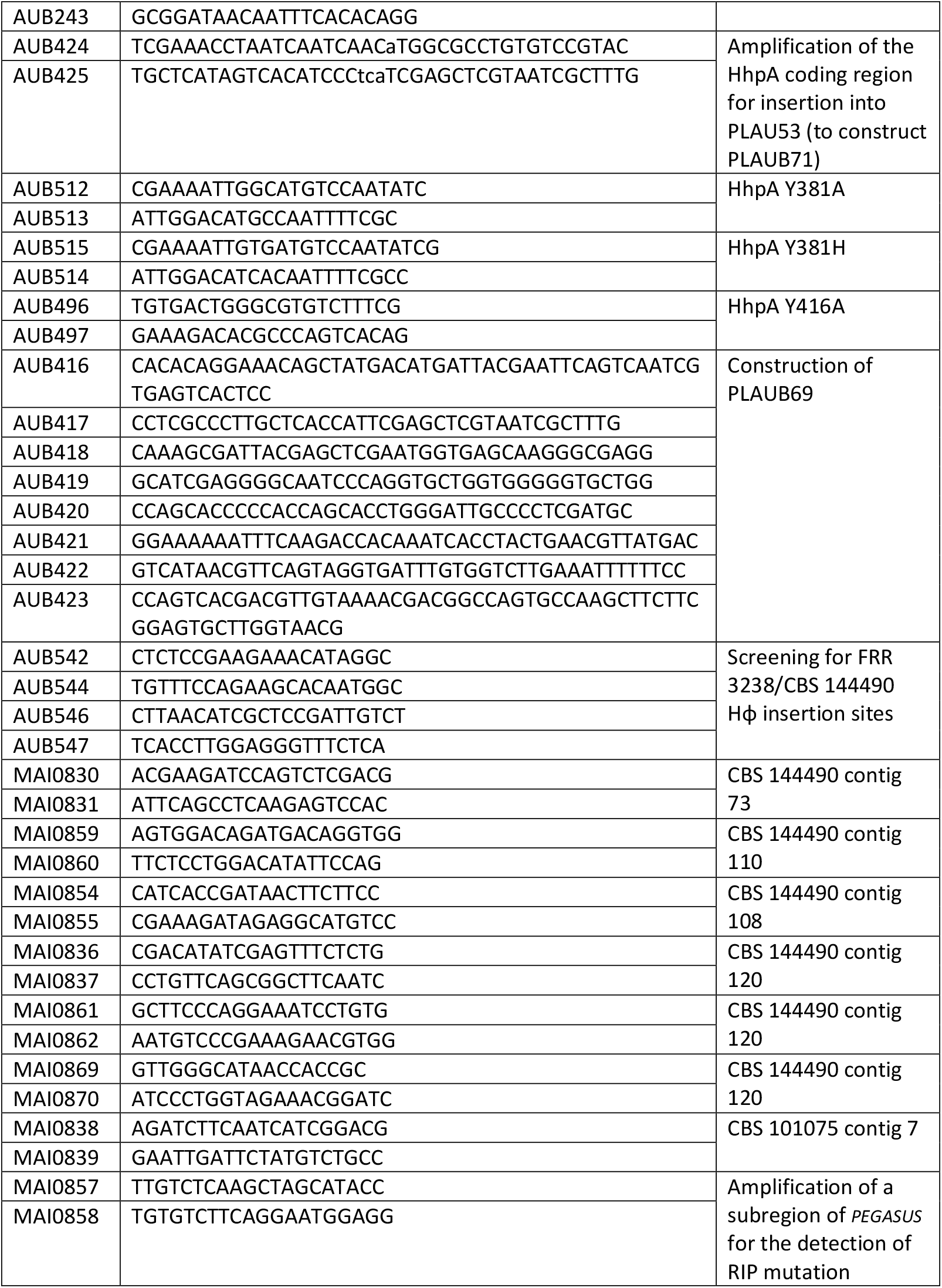
Oligonucleotides designed in this study:

**Table S7.**
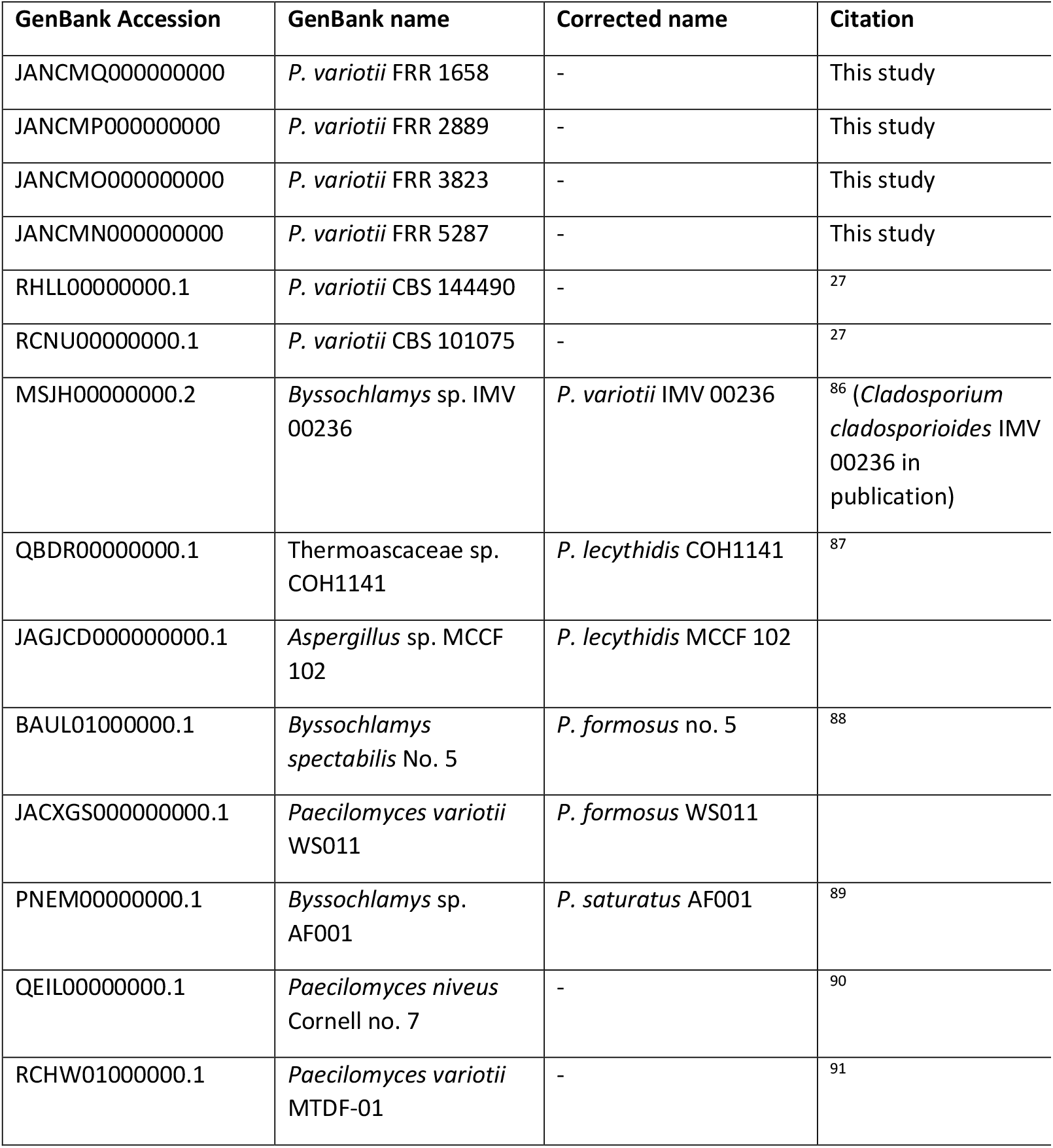
Sequenced *Paecilomyces* genomes.

**Table S8.** Illumina sequencing reads generated in this study.

| **Sequencing run** | **SRA accessions** |
| --- | --- |
| Pool of 13 spontaneously hygromycin resistant strains derived from the “double flanked” strain | SRR19913591 |
| Pool of several hundred *miniHϕ* spontaneous hygromycin resistant strains | SRR19913590 |
| RIP progeny 3 | SRR19913589 |
| RIP progeny 4 | SRR19913588 |
| RIP progeny 14 | SRR19913587 |
| RIP progeny 22 | SRR19913586 |
| RIP progeny 23 | SRR19913585 |

